# NSD1 inactivation defines an immune cold, DNA hypomethylated subtype in squamous cell carcinoma

**DOI:** 10.1101/178608

**Authors:** Kevin Brennan, June Ho Shin, Joshua K. Tay, Marcos Prunello, Andrew Gentles, John B. Sunwoo, Olivier Gevaert

## Abstract

Chromatin modifying enzymes are frequently mutated in cancer, resulting in a cascade of epigenetic deregulation. Recent reports indicate that inactivating mutations in the histone methyltransferase NSD1 define an intrinsic subtype of head and neck squamous cell carcinoma (HNSC) that features widespread DNA hypomethylation. Here, we describe a similar DNA hypomethylated subtype of lung squamous cell carcinoma (LUSC) that is enriched for both inactivating mutations and deletions in *NSD1*. The ‘NSD1 subtype’ of HNSC and LUSC are highly correlated at the DNA methylation and gene expression levels, with concordant DNA hypomethylation and overexpression of a strongly overlapping set of genes, a subset of which are also hypomethylated in Sotos syndrome, a congenital growth disorder caused by germline *NSD1* mutations. Further, the NSD1 subtype of HNSC displays an ‘immune cold’ phenotype characterized by low infiltration of tumor-associated leukocytes, particularly macrophages and CD8^+^ T cells, as well as low expression of genes encoding the immunotherapy target PD-1 immune checkpoint receptor and its ligands PD-L1 and PD-L2. Using an *in vivo* model, we demonstrate that NSD1 inactivation results in a reduction in the degree of T cell infiltration into the tumor microenvironment, implicating NSD1 as a tumor cell-intrinsic driver of an immune cold phenotype. These data have important implications for immunotherapy and reveal a general role of NSD1 in maintaining epigenetic repression.

## Introduction

Nuclear receptor binding SET domain protein 1 (*NSD1*) is frequently mutated in head and neck squamous cell carcinoma (HNSC) (1,2), the sixth most common cancer by incidence (3), and a leading cause of cancer-related death (4). NSD1 is also genetically or epigenetically deregulated (either inactivated or overexpressed) in several other cancer types (1,2,5–12).

NSD1 is best known as the causative gene for the congenital overgrowth disorder Sotos syndrome, which is associated with mildly increased cancer incidence (13–15). NSD1 is therefore among several epigenetic modifying enzymes (such as NSD2, DNMT3a, SETD2, EZH2) that represent causative genes for developmental growth disorders that are also frequently mutated in cancer (16).

*NSD1* is a SET-domain containing histone methylatransferase, which catalyzes methylation of histone 3 at lysine 36 (H3K36). Current evidence suggests that NSD1 catalyzes H3K36 dimethylation (H3K36me2) (17–19), though the precise epigenetic function of NSD1 (i.e. the H3K36 methylation states it catalyzes, its target genes and genomic loci, and the functional consequence of these marks) remains largely unknown.

Choufani et al. reported that germline *NSD1* mutations are associated with widespread perturbation (primarily loss) of DNA methylation (20), i.e., methylation of cytosine to form 5-methylcytosine at CpG dinucleotides. NSD1 is not thought to methylate DNA; therefore H3K36me (or other histone marks) catalyzed by NSD1 apparently regulate DNA methylation.

Inactivating mutations of *NSD1* also deregulate DNA methylation in HNSC, as we and others have described a HNSC subtype characterized by widespread DNA hypomethylation, that is strongly enriched for NSD1 mutations (2,19,21). We recently identified this ‘NSD1 subtype’ as one of five HNSC DNA methylation subtypes, using data from 528 HNSC patients from The Cancer Genome Atlas (TCGA) study (2,22). Papillon-Cavanagh et al. recently reported that a HNSC subtype featuring NSD1 mutations is defined by impairment of dimethylation (H3K36me2) and that NSD1 inactivation represents one of two mechanisms causing H3K36me2 impairment, the other being H3 K36M mutations (19). These findings reveal NSD1 inactivation as one mechanism underlying deregulation of DNA methylation, a major cause of abnormal gene expression in virtually all cancers (16).

Analysis of the gene expression profiles of these subtypes indicated striking inter-subtype differences in the profiles of both overall and cell type-specific tumor associated leukocytes (TALs). Tumors can exploit mechanisms of immune regulation to suppress infiltration of immune cells into the tumor microenvironment, thus avoiding anti-tumor immunity. There is a growing interest in identifying these mechanisms, which may be targeted using immunotherapies to restore innate anti-tumor immunity. Immunotherapies provide particular promise for metastatic HNSC; however they are only effective in a subset of individuals, and are associated with autoimmune side effects, therefore there is clinical need for biomarkers to identify patients that may be particularly sensitive. Current evidence indicates that ‘immune hot’ tumors, particularly those with greater numbers of infiltrating PD-1^+^ or? CD8^+^ T cells, are more responsive to immunotherapy (23), indicating that susceptibility to some immunotherapy approaches may vary between the HNSC subtypes.

Here, we follow up upon our subtyping analysis to describe the NSD1 subtype and report our identification of an epigenetically and transcriptionally similar NSD1 subtype occurring in lung squamous cell carcinoma (LUSC). We further investigated the immune profile of the HNSC NSD1 subtype and found that it represents an ‘immune cold’ subtype, with the lowest levels of overall and cell type-specific immune infiltrating lymphocytes among the five different HNSC tumor subtypes. We demonstrate that NSD1 inactivation induces immune cell exclusion from the tumor microenvironment using an *in vivo* mouse model of tumor immune infiltration, recapitulating the immune cold phenotype observed in the analysis of the TCGA data. These results may have important implications as a biomarker for the future selection of immune therapy-responsive patients.

## Methods and Materials

### Data processing

Preprocessed TCGA DNA methylation data (generated using the Illumina Infinium HumanMethylation450 and the HumanMethylation27 BeadChip arrays), gene expression data (generated by RNA sequencing), DNA copy number data (generated by microarray technology), and somatic point mutation data (generated by genome sequencing) were downloaded using the Firehose pipeline (version 2014071500 for gene expression and version 2014041600 for all other data sets) (24). Copy number was called using GISTIC2.0. RNA-Seq data was processed using RSEM. Preprocessing for these data sets was done according to the Firehose pipelines described elsewhere (24). Mutation data was accessed as Mutation Analysis reports, generated using MutSig CV v2.0 (25). Mutations predicted as silent by MutSig CV were removed. Additional data preprocessing of gene expression and DNA methylation data was done as follows: Genes and patients with more than 10% missing values for gene expression, and more than 20% missing values for DNA methylation, were removed. All remaining missing values were estimated using KNN impute (26). Batch correction was done using Combat (27).

### Classification of abnormally methylated genes

To reduce multiple testing of highly correlated CpG probes, probes for each gene were clustered using hierarchical clustering with complete linkage, and mean methylation (beta-value) was calculated for each CpG cluster. MethylMix was applied to CpG cluster data to systematically identify regional CpG clusters that are abnormally methylated in cancer versus normal tissue, where DNA methylation is inversely associated with RNA expression of the corresponding gene, using beta-mixture models, as previously described (28). For each gene (CpG cluster), MethylMix ascribes either normal or abnormal (hypomethylated or hypermethylated) DNA methylation states to each patient. For LUSC, 370 patients had DNA methylation data generated using the Illumina 450k array, while 133 patients had methylation data measured using the Illumina 27k array. To maximize the methylation data in terms of either patient numbers or genomic coverage, depending on the application, MethylMix was applied twice: first to all 503 patients, using data for CpG probes that were shared between the 450k and 27k array platforms (n=23,362 probes), and then to separately the 370 patients with 450k array data (n=395,772). For HNSC, all 528 patients had DNA methylation data generated using the Illumina 450k array.

### Consensus clustering of abnormally methylated genes

Consensus clustering was applied to MethylMix output data, i.e. methylation state data, for cancer patients, to identify robust patient clusters (Putative subtypes). Consensus clustering was performed using the ConsensusClusterPlus R package (29), with 1000 rounds of k-means clustering and a maximum of k=10 clusters. Selection of the best number of clusters was based on visual inspection ConsensusClusterPlus output plots. For HNSC, subtypes are as previously described (22). For LUSC, consensus clustering was applied to MethylMix output data for all 503 patients, in order to maximize the number of patients with both mutation and DNA methylation data.

### Identification of genes associated with NSD1 subtypes

SAM analysis (30) was used to identify genes that were overexpressed and underexpressed NSD1 subtypes relative to all other patients. SAM analysis was also used to identify genes (CpG clusters) that were either hypermethylated or hypomethylated within the NSD1 subtypes, using mean methylation for each CpG cluster. For LUSC, SAM analysis was applied only to DNA methylation data for he 370 patients with 450k array data (Excluding patients wit 27k data), to maximize genome coverage.

### Centroid-based classification of LUSC patients to the HNSC NSD1 subtype

Prediction Analysis of Microarrays (PAM) (31) was used to develop a DNA methylation classifier to predict the HNSC NSD1 subtype, and to classify LUSC patients that are most similar to the HNSC NSD1 subtype at the level of DNA methylation. Briefly, PAM uses a nearest shrunken centroids method to assign the class of each LUSC patients based on the squared distance of the DNA methylation profile for that individual to the centroids of known class groups (i.e. HNSC patients within, or not within the NSD1 subtype).

We applied PAM to DNA methylation data for all 10,818 CpG sites within all gene regions that were abnormally methylated (Hypomethylated or hypermethylated) in HNSC, identified using MethylMix (28), as previously reported (22). PAM analysis uses Shrinkage to select the optimum number of CpG probes for class prediction, such that the model selects only a subset of CpG probes to develop the centroids. We first used PAM in combination with 10-fold cross validation to determine the ability of the DNA methylation data to predict the NSD1 subtype within TCGA data. For each fold of cross validation, the PAM model was trained on 90% of patients and assigned class probability for belonging to the NSD1 subtype to the each of the remaining 10% of patients based on the distance of the patient to its closest centroid. We used the Area under the ROC curve (AUC) to evaluate the performance of the model in accurately predicting the class of samples. We then applied this DNA methylation classifier signature to 365 TCGA LUSC patients (All patients with 450k array data) to classify them into either a ‘HNSC NSD1 subtype’ class or the ‘HNSC other subtype’ class. We only used classification results when probabilities were >60% or <40%, excluding low confidence assignments for one borderline individual from analyses.

### Inference of tumor associated leukocyte levels

CIBERSORT was applied to gene expression (RNA-Seq) data to infer the levels of specific TAL types, as previously described (32,33). Only patients for with estimation p-values less than 0.05 (Indicating high confidence TAL estimation) were included in downstream analyses.

### Inference of infiltrating T cells using a T cell gene expression signature

Mean expression of a set of 13 T cell transcripts (*CD8A*, *CCL2*, *CCL3*, *CCL4*, *CXCL9*, *CXCL10*, *ICOS*, *GZMK*, *IRF1*, *HLA-DMA*, *HLA-DMB*, *HLA-DOA*, *HLA-DOB*), across all 13 genes, was used as a method of inferring relative T cell levels, as previously described (34). This T cell score was strongly correlated with expression of CD8+ T cell expressed *PDCD1* and negatively associated with expression of *EPCAM*, a marker of epithelial tumor purity (Low stromal/immune content) (35) (Supplementary figure 1).

### Processing copy number data (For the GSE33232 studycessing copy number data (For the GSE33232 study (36) cohcohort)

Raw CEL signal intensity files (Generated using the Affymetrix Genome-Wide Human SNP 6.0 Array) were processed with Affymetrix power tools and BIRDSUITE 1.5.5 (37). Segmented copy-number calls were log2 transformed and further processed with GISTIC (38) using an amplification and deletion threshold of 0.1. Samples with *NSD1* copy number calls meeting the GISTIC threshold and designated at least −1 or +1 were considered to have *NSD1* deletions and amplifications, respectively.

### Mice and cell lines

NSG mice (NOD-scid IL2Rgamma^null^, 6-12 weeks old) on a C57BL/6 background were a kind gift from Dr. Ravi Majeti (Stanford University) and were bred in our animal facility under pathogen-free conditions. All animal procedures were performed in accordance with protocols approved by the Administrative Panel on Laboratory Animal Care at Stanford University (Stanford, CA).

The human HNSC cell lines PCI-13 was a gift of Suzanne Gollin at the University of Pittsburgh. The UM-SCC-6 cell line was obtained from the University of Michigan. The FaDu cell line was obtained from ATCC. Cells were maintained in complete DMEM:F12 medium (DMEM:F12 1:1 with 10% heat-inactivated FBS [Omega Scientific], 100 IU/ml penicillin and 100 μg/ml streptomycin [Gibco, Invitrogen, CA]). The 293T cell line was obtained from ATCC and maintained in complete DMEM medium. Culture medium was changed every 2–3 days depending on cell density, and subculture was conducted when confluence was reached.

### Lentiviral shRNA transduction

For the production of the lentiviral particles, 293T cells were transfected using Lipofectamin2000 (Invitrogen) with the packaging plasmid pCMVR8.74 (Addgene), the envelope plasmid pCMV-VSVG, and the lentiviral construct containing the human NSD1 shRNA (pGIPz lentiviral vector, Dharmacon GE Life Sciences). Cell culture medium was changed 16 hours after the transfection and virus supernatants were collected 24 and 48 hours after the media change. Immediately after supernatant collection, the viral particles were concentrated by polyethylene glycol precipitation with PEGit solution (SBI Bioscience), according to the manufacturer’s protocol. The lentiviral pellets were then resuspended in ice-cold PBS. For the lentiviral transduction of the cell lines, cells were plated at the appropriate concentration (1x10^5 cells per 6 well plates). Then, the lentiviral particles were added to the cell cultures at a multiplicity of infection (MOI) of 1 transducing Unit per cell. Polybrene (8ug/ml) was also added to enhance the lentiviral transduction efficiency. Medium was changed 24 hours after viral infection. All transfected cells were purified by FACS sorting for GFP^+^ cells and expanded for the experiments.

### RNA extraction and chemokine gene expression array

RNA was extracted with the RNeasy mini kit (QIAGEN), and cDNA made with the Maxima First Strand cDNA Kit (ThermoFisher Scientific). For chemokine gene expression assessment, a TaqMan human chemokine and cytokine array was purchased from ThermoFisher Scientific and was used per the manufacturers protocol. The amplified cDNA was diluted with nuclease-free water and added to the TaqMan^®^ Gene Expression Master Mix (ThermoFisher Scienticfic). Then, 20 µl of the experimental cocktail was added to each well of the TaqMan™ Array Human Chemokines (ThermoFisher Scienticfic, CA). Real-Time PCR was performed on the 7900HT Fast Real-Time PCR System (Applied Biosystems, CA) with the following thermal profile: segment 1 - 1 cycle: 95°C for 10 minutes, segment 2 - 40 cycles: 95°C for 15 seconds followed by 60°C for 1 minute, segment 3 (dissociation curve) - 95°C for 1 minute, 55°C 30 seconds, and 95°C for 30 seconds.

### In vivo tumor infiltration assay and flow cytometry

Control and NSD1 shRNA knockdown HNSC cells (1x10^6) were injected into the subcutaneous compartment of the flanks of NSG mice. In each mouse, one flank was injected with control cells and the other with an equal number of NSD1 knockdown cells. After tumors were established (5 mm diameter), 100 × 10^6^ Ficoll-purified human PBMCs per mouse were injected via tail vein. After 10 days, tumors were dissociated, and tumor-infiltrating T cells (CD45^+^CD3^+^) were quantified by FACS. Human PBMCs were obtained from healthy volunteers at the Stanford Blood Center (Palo Alto, CA) and prepared by Ficoll gradient centrifugation (GE Healthcare, Piscataway, NJ, USA). For tumor digestion, tumors were isolated/minced and digested in 300 U/mL collagenase and 100 U/mL hyaluronidase (StemCell Technologies) in culture media; DMEM/F-12 medium (Corning) with 10% FBS, and 1% penicillin-streptomycin-amphotericin B (ThermoFisher Scienticfic). The tumor digest was pipetted every 15 minutes and incubated at 37°C for 3 hours, until a single-cell suspension was obtained. The dissociated cells were spun down and resuspended in Trypsin/EDTA (StemCell Technologies) for 5 minutes, then further dissociated with 5 U/mL dispase (StemCell Technologies) and 0.1 mg/mL DNase I (StemCell Technologies) for 1 minute. Cells were filtered through a 40-mm cell strainer and erythrocytes were lysed with ACK lysing buffer (Lonza) prior to antibody staining and FACS. The dissociated cells were resuspended in ice cold FACS solution (PBS supplemented with 2% fetal calf serum and 1% penicillin-streptomycin) and stained with PerCP-Cy5.5-anti-human CD3, APC-anti-human CD45 (BioLegend, CA) according to the manufacturer’s protocols. DAPI (1 µg/mL) was added to all the tubes prior to filtering through a 70 µm membrane. Labeled cells were analyzed on a FACSAria III (BD Biosciences).

## Results

### Association of NSD1 mutations and deletions with a DNA hypomethylated HNSC subtype

We recently described a HNSC subtype featuring widespread DNA hypomethylation cooccurring with NSD1 mutations using MethylMix (21,22). Of 2,602 genes found to be abnormally methylated in HNSC relative to normal tissue overall (22), 1127 were significantly hypomethylated, and 102 hypermethylated, in the *NSD1* subtype relative to other HNSC subtypes combined (Supplementary table 1). Fifty-seven percent (24/42) of patients within this HNSC subtype had *NSD1* mutations, compared with 2-8% patients within the other subtypes. This subtype included all five patients with ‘high-level’ somatic deletions called by GISTIC 2.0 (38), as well as enrichment of ‘low-level’ deletions. NSD1 deletions were significantly enriched among patients with NSD1 point mutations, as 21/33 (64%) of patients with NSD1 mutations had deletions, compared with 99/269 (0.34) of patients without mutations. However, mutations and deletions were each independently associated with both *NSD1* RNA expression (Supplementary figure 2a) and mean DNA methylation across all abnormally methylated genes (Supplementary figure 2b). Lowest NSD1 expression and mean methylation occurred in patients with high-level likely biallelic deletions but without mutations, and in patients with both NSD1 mutations and deletions, suggesting that tumors undergo positive selection for loss of both alleles, resulting in extreme hypomethylation. Moreover, patients with low-level deletions had significantly lower mean DNA methylation in patients with and without NSD1 mutations, indicating that NSD1 deletions generally impair DNA methylation.

### Identification of a hypomethylated, NSD1 inactivated subtype of lung squamous cell carcinoma

We investigated the possibility that NSD1 mutations affect DNA methylation in other cancers, focusing on cancers for which there were at least ten patients with *NSD1* mutations and accompanying DNA methylation data within TCGA data. These included LUSC, uterine corpus endometrial carcinoma (UCEC), and breast carcinoma (BRCA). LUSC was the only of these cancers in which NSD1 mutations were significantly associated with DNA hypomethylation (p=0.001) (Supplementary figure 3).

To investigate whether NSD1 inactivation occurred within a hypomethylated subtype of LUSC, as is the case for HNSC, we performed consensus clustering of 503 LUSC patients based on their profiles of abnormally methylated genes identified using MethylMix (28), the method that revealed the HNSC NSD1 subtype (21,22). This resulted in 3,025 abnormally methylated genes, consensus cluster of which revealed six subtypes. One of these subtypes had a significantly elevated number of hypomethylated genes (Figure 1). This subtype included six of ten LUSC patients with NSD1 mutations, representing 17% of patients in this subtype (p=0.005). This subtype was also enriched for NSD1 deletions, as 88/104 (84%) of patients within this subtype had deletions compared with 31-74% patients within other subtypes (p=0.001). NSD1 RNA expression and DNA methylation displayed the same inverse trend with mutations and deletions, as seen in HNSC (Supplementary figures 2).

**Figure 1:**
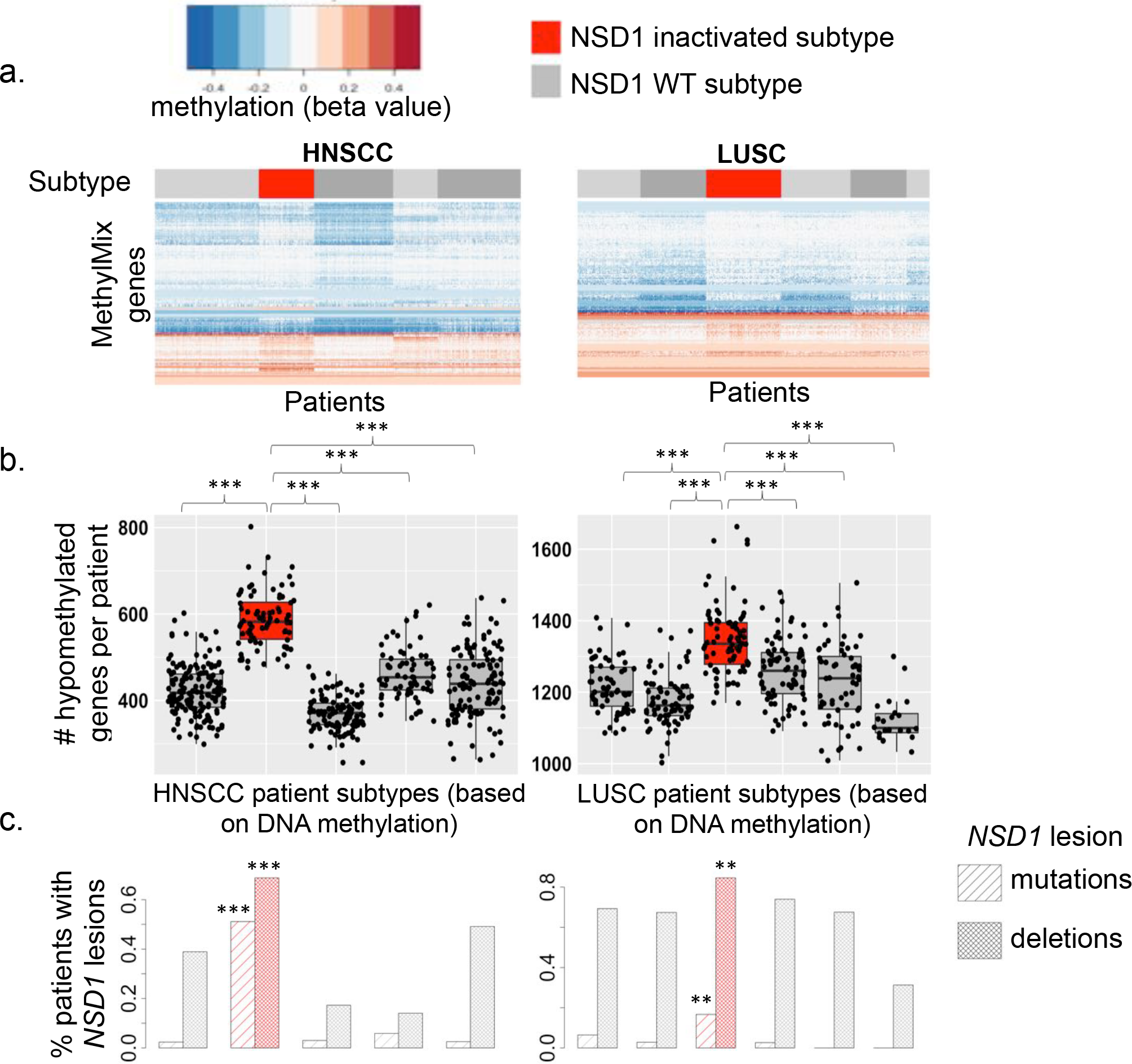
Identification of NSD1 inactivated subtypes of squamous cell carcinomas featuring epigenetic de-repression of developmental oncogenes: a) Heatmaps illustrate five subtypes of head and neck squamous cell carcinoma (HNSC, n=528 patients) and six subtypes of lung squamous cell carcinoma (LUSC, n=502 patients) within TCGA studies, identified by consensus clustering of patients according to their profiles of abnormally methylated genes, subsequent to identification of these abnormally methylated genes by applying MethylMix to integrate DNA methylation and gene expression data. Red bars demarcate hypomethylated NSD1 subtypes, while light and dark grey bars demarcate other subtypes. b) The average number of genes hypomethylated per patient (in tumor relative to normal tissue) was significantly higher in NSD1 subtypes (red) than each other subtype (grey) in both HNSC and LUSC. c) Percentages of patients within each subtype that have NSD1 mutations (Striped bars) and NSD1 deletions (Solid bars) in HNSC and LUSC. Asterisks indicate the significance of enrichment of NSD1 mutations or deletions within the NSD1 subtype (red) compared with patients in all other subtypes (Pearson’s chi-squared test).

The DNA methylation profiles of the HNSC and LUSC NSD1 subtypes are strongly concordant, illustrated by a correlation matrix heatmap indicating pairwise correlations between each HNSC patients and LUSC patients (Figure 2a). Previous investigations have identified concordance between HNSC and LUSC gene expression subtypes, using centroid predictor based approaches (2,39). We used a similar method, PAM analysis (31), to classify those LUSC patients that are similar to the HNSC NSD1 subtype, and HNSC patients that are similar to the LUSC NSD1 subtype, based on their DNA methylation profiles. We first trained PAM models to classify the NSD1 subtype and tested their accuracy using internal 10-fold cross validation, within HNSC and LUSC separately. These PAM models for HNSC and LUSC could classify NSD1 subtype patients with areas under the receiver-operating curve (AUC) of 0.997 (95% CI: 0.991-1), and 0. 86 (0.81-0.90), respectively. The AUC for the HNSC PAM model remained high (0.96 (95% CI: 0.94-0.99)) when the number of CpG sites used for class prediction was reduced to just five, indicating that it would be possible to identify the HNSC NSD1 subtype using a minimal CpG panel biomarker.

**Figure 2.**
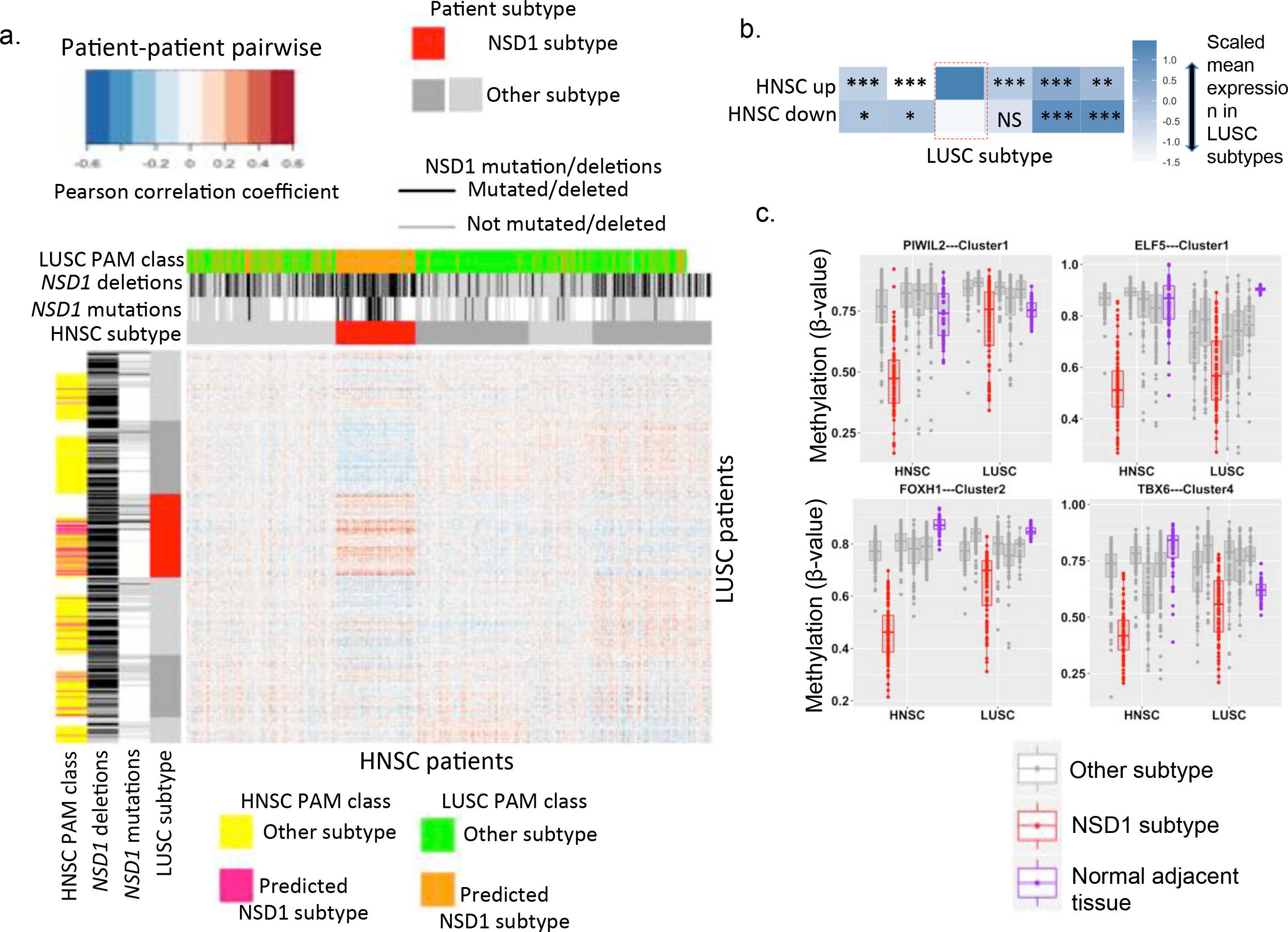
Concordant DNA methylation and gene profiles between HNSC and LUSC subtypes: a) Heatmap illustrating of a correlation matrix illustrating pair-wise correlations between 528 HNSC patient and 503 LUSC samples, based on DNA methylation data (Pearson’s correlation coefficients). DNA methylation data included 621 probes, representing all CpG sites that were within HNSC MethylMix genes, i.e., gene regions that were abnormally methylated in HNSC tumor relative to normal tissue, that were available for all patients (Measured on both Illumina 27k and 450k arrays). Subtype sidebars indicate HNSC and LUSC subtypes (NSD1 subtypes illustrated in red, other subtypes grey). NSD1 mutation and deletion sidebars indicate patients with NSD1 mutations or deletions (black), absence of NSD1 mutations or deletions (grey), or missing data (white). The ‘NSD1 PAM clas’ sidebar indicates PAM analysis class predictions for LUSC patients, using a model to identify patients that are more similar to the HNSC NSD1 subtype (pink) or HNSC patients not within the NSD1 subtype (yellow), based on their DNA methylation profiles. b) Scaled mean RNA expression in LUSC DNA methylation subtypes of genes that were upregulated (HNSC up) and downregulated (HNSC down) in the HNSC NSD1 subtype. Asterisks indicate the significance of differential mean expression between the NSD1 LUSC subtype (Red box) and each other subtype (Wilcoxon rank sum test): NS Not significant, * P<0.05, ** P<0.01, *** P<0.001. c) DNA methylation of development-related transcription factor genes, in normal tumor-adjacent tissue (purple), and in tumor of patients within NSD1 subtypes (red) or other subtypes (grey), in HNSC and LUSC.

We validated the HNSC PAM model by applying it to an independent set of 44 primary HNSCs, for which methylation, RNA expression and copy number data was available (GSE33232) (36). Six (14%) of these HNSCs that were classified as the NSD1 subtype. These predicted NSD1 subtype patients had significantly lower *NSD1* RNA expression (p=0.014, Supplementary figure 4). Interestingly, *NSD1* RNA expression was negatively correlated with methylation of genes that were hypermethylated in the HNSC subtype, as well as positively associated with genes that were hypomethylated, confirming that NSD1 inactivation causes DNA hypermethylation as well as hypomethylation. Both patients with *NSD1* deletions were within the group predicted as the NSD1 subtype. This indicated that the HNSC NSD1 subtype PAM model could classify NSD1 subtype patients in external data sets.

We next applied the HNSC PAM model to LUSC patients, and found that 58/365 (16%) of patients were assigned to the HNSC NSD1 subtype class, of which 35 (60%) were within the LUSC NSD1 subtype, representing a strong enrichment (p=5.6e-15) (Figure 2a). Conversely, when we applied the LUSC PAM model to HNSC, 165/527 (31%) of patients were assigned to the LUSC NSD1 subtype class, of which 79 (48%) were within the HNSC NSD1 subtype (p<2.2e-16) (Figure 2a). This confirmed the similarity of the HNSC and LUSC NSD1 subtypes at the DNA methylation level.

The HNSC and LUSC NSD1 subtypes were also concordant at the transcriptional level, as mean expression of genes upregulated and downregulated in the HNSC NSD1 subtype were upregulated and downregulated, respectively, in the LUSC NSD1 subtype, compared with each other subtype (Figure 2b). The molecular similarity of the HNSC and LUSC NSD1 subtypes was primarily driven by DNA hypomethylation concordant with transcriptional upregulation, as 178/867 (20%) genes that were significantly overexpressed within the HNSC NSD1 subtype were also overexpressed within the LUSC NSD1 subtype (Supplementary table 1), while 37/722 (5%) genes underexpressed with the HNSC NSD1 subtype were underexpressed within the LUSC NSD1 subtype (Supplementary table 2).

Intriguingly, among the genes hypomethylated and overexpressed in both HNSC and LUSC NSD1 subtypes were transcription factors that are normally expressed specifically in germline tissues or during development, for example, *PIWIL2* (40,41), *ELF5* (42), *TBX6* (43) and *FOXH1* (44). These genes were highly methylated in adjacent normal tissues, but hypomethylated at functional gene regions, often promoter CpG islands (Supplementary Table 1), specifically within NSD1 subtypes.

### The cancer NSD1 DNA hypomethylation signature overlaps with the Sotos syndrome hypomethylation signature

Using a reported set of CpG sites that are abnormally methylated in Sotos syndrome (20), we investigated the possibility that a shared set of genes is epigenetically deregulated by NSD1 in different diseases. Of 49 CpG probes hypermethylated in Sotos syndrome, none were hypermethylated in either HNSC or LUSC. However, of 7,038 probes hypomethylated in Sotos syndrome, 117 were hypomethylated in the HNSC NSD1 subtype, and 161 were hypomethylated in the LUSC NSD1 subtypes, with 54 hypomethylated probes within 31 unique genes overlapping between Sotos syndrome, HNSC and LUSC (p<2.2e-16) (Supplementary table 1). For each cancer patient, we calculated the percentage of hypomethylated probes that overlapped with the Sotos signature and determined whether the extent of overlap varied between cancers with and without *NSD1* lesions (Mutations or deletions). This revealed a trend whereby overlap with the Sotos syndrome hypomethylation signature increased incrementally with an increasing number of *NSD1* lesions in both HNSC and LUSC (Supplementary figure 5), suggesting that NSD1 inactivation induces hypomethylation of a discrete set of genes across disease states and tissue types.

### NSD1 inactivation is associated with an immune cold phenotype in HNSC

We recently reported that levels of tumor associated leukocytes (TALs), inferred from gene expression data using the CIBERSORT algorithm (32,33), varied between HNSC DNA methylation subtypes (22) (Figure 3a). The NSD1 subtype displayed an ‘immune cold’ subtype, displaying the lowest overall TAL levels as well as the lowest levels of specific TAL types including pro-inflammatory M1 macrophages, CD8^+^ cytotoxic T cells and resting CD4^+^ memory T cells, while plasma cells were highest within the NSD1 subtype.

**Figure 3:**
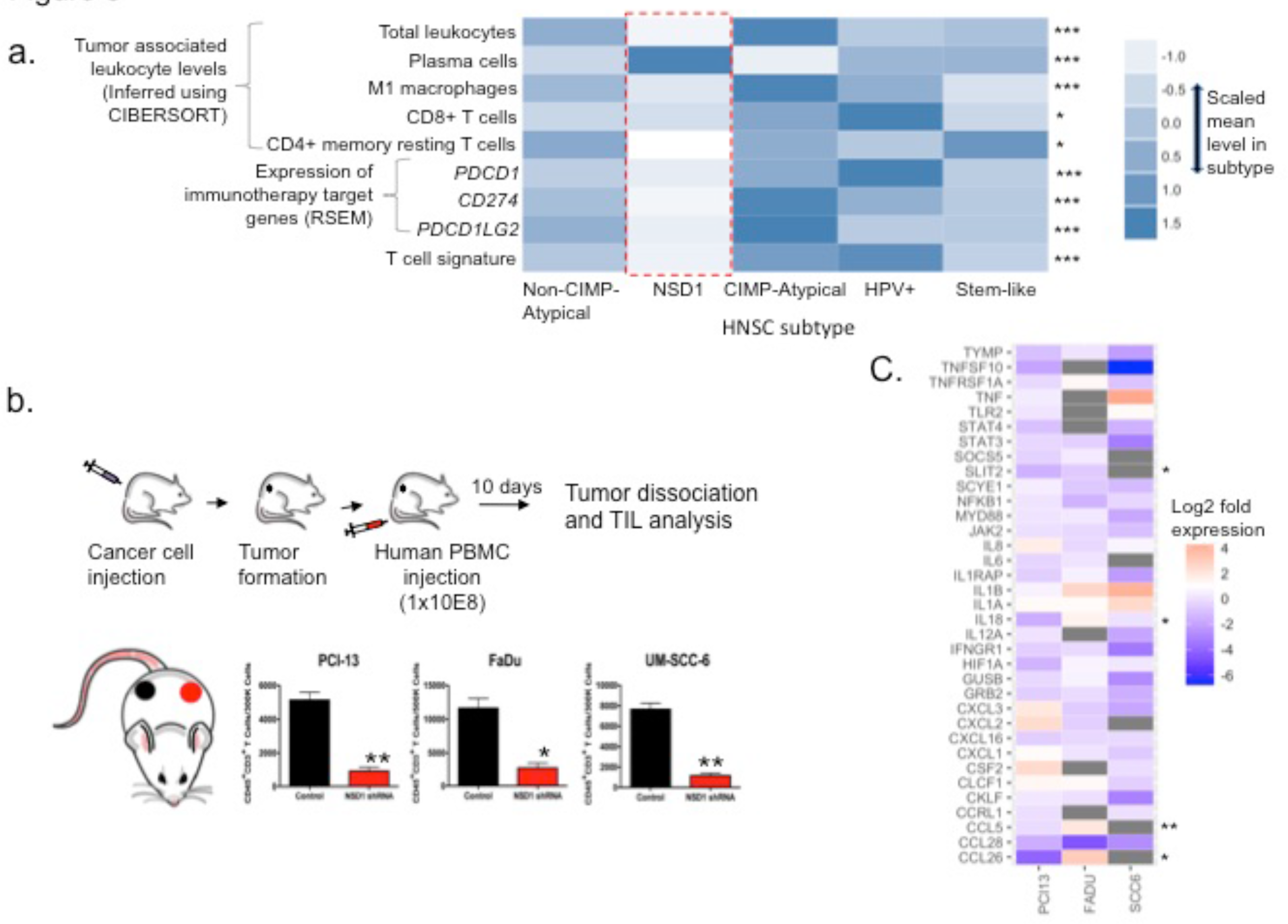
NSD1 inactivation is associated with immune cell exclusion from the tumor microenvironment in HNSC: a) Compared with other HNSC subtypes, the NSD1 subtype (red box) displayed significantly lower mean signature levels of overall tumor associated leukocytes (TALs), and specific TAL types including M1 tumor associated macrophages (TAMs), CD8+ cytotoxic T cells, and CD4+ memory T cells (All inferred using CIBERSORT(32)). The NSD1 subtype had the low mean RNA expression of immunotherapy-relevant genes, including CD274 (PD-L1), PDCD1 (PD-1) and PDCD1LG2 (PD-L2), and a lower mean level of T cell signature based on expression of 13 T cell transcripts. B) Control and NSD1 shRNA knockdown HNSC cells (1x10^6) were injected into the subcutaneous compartments of the flanks of NOD-scid IL2Rgamma^null^ (NSG) mice. In each mouse, one flank was injected with control cells (black) and the other with NSD1 knockdown cells (red). After tumors were established (5 mm diameter), 100 × 10^6^ Ficoll-purified human PBMCs per mouse were injected via tail vein. After 10 days, tumors were dissociated, and tumor-infiltrating T cells (CD45^+^CD3^+^) were quantified by FACS. Cohorts were n=5 per set of control and knock-down cell line, as indicated. *P<0.05; **P<0.005 (paired two-tailed Students t-test, error bars represent S.D.). *C) NSD1 knockdown in HNSC results in the decreased expression of multiple chemokine genes*. Control and NSD1 shRNA knockdown HNSC cells were assessed for the expression of chemokine and chemokine-related genes using a qRT-PCR array. Log2 fold expression of of 35 chemokine-related genes upon NSD1 knockdown (Relative expression NSD1-shRNA/Control) in three established HNSC cell lines (PCI13, FADU, SCC6). Log2 fold expression is indicated by a color gradient, with NA values indicated in grey. Asterisks indicate genes that were upregulated* or downregulated** in the NSD1 subtype (relative to other subtypes) in the TCGA study.

Interestingly, the NSD1 subtype displayed low RNA expression of genes of relevance to immunotherapy, including the immune checkpoint receptor *PDCD1* (encoding PD-1), as well as its ligands *CD274* (encoding PD-L1) and *PDCD1LG2* (encoding PD-L2) (Figure 3a).

It is widely understood that PD-1 expressed on CD8^+^ T cells binds PD-L1 and/or PD-L2 expressed on tumor cells or other cells within the microenvironment, resulting in suppression of anti-tumor immune response. A recent report indicates that PD-1 is also expressed on tumor associated macrophages (TAMs), that the PD-1/PDL1 checkpoint inhibits phagocytosis of tumor cells by TAMs, and that PD-1-PDL1 blockade immunotherapy functions through reactivation of TAMs as well as CD8^+^ T cells (45). The authors reported that PD-1 is particularly expressed on alternatively activated M2, rather than classically activated M1 TAMs, based on cell surface protein markers. The co-occurrence of low *PDCD1* expression and M1, but not M2 TAM levels in the NSD1 subtype led us to hypothesize that PD-1 expression may actually be associated with M1 TAM levels; therefore, we investigated the correlation of PD-1 expression with different TAM fractions inferred by CIBERSORT, across 28 TCGA cancer types. Indeed, *PDCD1* expression was positively correlated with M1 macrophage and CD8^+^ T cells (Supplementary figure 6). This postulates that M1 TAMs represent the TAM fraction that express PD-1 and are susceptible to reactivation by immunotherapy. Consistent with recent reports that TAMs are reprogrammed to express PD-L1 (46–48), M1 macrophage levels were also specifically correlated with expression of *CD274* and *PDCD1LG2* (Supplementary figure 6). Given that both M1 TAMs and CD8^+^ T cells, as well as that *PDCD1*, *CD274* and, *PDCD1LG2* are lowest within the NSD1 HNSC subtype, we speculate that the NSD1 subtype is particularly immune evasive, and may be highly resistant to immunotherapy.

Using NSD1 RNA expression as a measure of NSD1 proficiency, we next validated the correlation of NSD1 expression with tumor infiltrating T cell levels in three independent primary HNSC population data sets, including the aforementioned GSE33232 data set and two additional datasets: GSE65858 (n=253) (49) and GSE39366 (n=138) (39). As a marker of T cell infiltration, we used a T cell signature based on mean expression of 13 T cell transcripts, previously employed elsewhere (34). NSD1 RNA expression was positively correlated with T cell levels in all three independent patient cohorts, although the correlation was not statistically significance in the smallest (GSE33232) data set (Supplementary figure 7). This indicates that NSD1 expression represents a reproducible marker of T cell infiltration in HNSC.

### Knockdown of NSD1 in HNSC results in immune cell exclusion from the tumor microenvironment

To assess a potential functional role of NSD1 inactivation in the exclusion of immune cells from the tumor microenvironment, we inhibited the expression of NSD1 by shRNA transduction in three established HNSC cell lines, PCI-13, FaDu, and UM-SCC-6. Matched sets of control and NSD1 knockdown cells were used to establish tumors in opposite flanks of immunodeficient NOD-scid IL2Rgamma^null^ (NSG) mice (Figure 3b). Once tumors formed, human peripheral blood mononuclear cells (PBMCs) were injected intravenously, and the degree of T cell infiltration into the tumors was assessed by dissociation of the tumors and analysis of infiltrating T cell levels by flow cytometry. There was a significantly lower number of T cells in the NSD1 knockdown tumors compared to the control transduced tumors established from the three sets of cell lines. This points to a functional role of NSD1 inactivation in the exclusion of immune cells from the tumor microenvironment and is consistent with our observations of a correlation between NSD1 expression and T cell infiltration (Figure 3a and Supplementary figure 7). To begin to understand how NSD1 inactivation may be affecting T cell infiltration, we compared the expression of an array of chemokine genes in the control cell lines to matched NSD1 knockdown cell lines. The expression of multiple key chemokines important for immune cell recruitment was downregulated in the NSD1 knockdown cells (Figure 3c), consistent with the reduction in the number of infiltrating T cells in NSD1 knockdown tumors. Thus, our data support a role of NSD1 as a tumor cell-intrinsic determinant of T cell infiltration into the tumor microenvironment.

## Discussion

Here we have described a hypomethylated, immune cold subtype of HNSC that is enriched for NSD1 mutations and somatic deletions, as well as a molecularly similar subtype in LUSC.

Our analysis indicates that both NSD1 mutations and deletions contribute significantly and independently to genome-wide deregulation and DNA methylation and transcription in a significant proportion of HNSCs and LUSCs. Indeed, our findings suggest that the most pronounced hypomethylation occurs due to biallelic loss of NSD1 at the transcriptional level, associated with combined mutations and deletions. Detailed genetic studies will be required to definitively characterize pathogenic lesions.

The NSD1 subtypes of HNSC and LUSC are characterized by DNA hypomethylation of many genes, concurrent with hypermethylation of smaller set, resulting a net loss of ‘global’ DNA methylation. This indicates that NSD1 inactivation does not simply preclude DNA methylation, but alters its distribution, and implies a complex role of NSD1 in locus-specific epigenetic regulation.

An emerging consequence of cancer DNA hypomethylation is loss of epigenetic repression of developmental or germline tissue-specific genes (71,72). This occurs in NSD1 squamous cell carcinoma subtypes, where concurrent hypomethylation and overexpression of developmental transcription factors such as *PIWIL2* (71), *ELF5*,(42) *TBX6*, (43) and *FOXH1* (44) occurs. Such ectopically expressed genes may play oncogenic roles, as *PIWIL2* and *ELF5* represent epigenetically-regulated oncogenes that promote oncogenic transcriptional networks in lung and other cancers (40,41,73–75). *PIWIL2* is among 31 genes that were hypomethylated in HNSC, LUSC, and Sotos syndrome, raising the intriguing possibility that genes and pathways that are responsible for overgrowth and cancer susceptibility in Sotos syndrome also promote growth in sporadic cancers.

NSD1 inactivation likely deregulates DNA methylation indirectly through alteration of underlying chromatin marks, as is the case of mutations in *SETD2* and MLL enzymes (50,51). NSD1 inactivation could deregulate DNA methylation by impairing H3K36 trimethylation (H3K36me3), a mark that regulates DNA methylation (52–54), as H3K36me1 and H3K36me2, the presumed methyltransferase products of NSD1 (17–19), represent substrates for conversion to H3K36me3 by SETD2 (55,56). Consistently, some (10,11,55), though not all (19) studies have found that NSD1 inactivation results in H3K36me3 loss. Interestingly, *SETD2* mutations, resulting in redistribution of H3K36me3, cause DNA hypermethylation at gene bodies in renal cell carcinoma (52), contrasting with widespread promoter hypomethylation in NSD1-inactivated cancers.

It is generally understood that HNSC and LUSC are molecular similar, as these cancer types tend to cluster together in pan-cancer unsupervised clustering analyses (21,67,68). Our analysis revealed a particularly striking correlation of the NSD1 subtypes between these two tumor types, revealing NSD1 inactivation as a driver of this novel molecular pan-cancer group. The defining feature of the NSD1 subtypes is likely to be loss of H3K36 methylation, resulting in altered DNA methylation and transcription. NSD1 genetic lesions represent one mechanism underlying impaired H3K36me; however, other mechanisms, such as H3K36 M mutations (19) or those that impair NSD1 at the protein level, may account for H3K36me loss within the NSD1 wild type cancers within these subtypes.

Inference of TAL levels based on gene expression data revealed that the HNSC NSD1 subtype displays an ‘immune cold’ phenotype characterized by lower levels of overall TALs, and M1 TAMs, CD8+ T cells and resting CD4 memory T cells in particular. The correlation of *NSD1* RNA expression with a T cell signature was consistent in three independent patient cohorts. Lower T cell levels within the NSD1 subtype are particularly clinically interesting, as T cell levels (particularly CD8^+^ T cells) represent markers of anti-cancer immune response that are associated with favorable prognosis in HNSC and other solid cancers (34,57–62). Thus, our findings may have important implications for the future selection of immune therapy-responsive patients.

There is a growing interest in identifying the determinants of tumor immune infiltration, particularly of immune cell factions that mediate anti-tumor immunity, such as CD8+ T cells and macrophages. Tumors can repress anti-tumor immune response by exploiting mechanisms of immune regulation, that normally function to prevent autoimmunity, such as by expressing ligands that activate immune checkpoints or by modulating expression of immune cells within the tumor microenvironment.

We have found intriguing evidence that NSD1 inactivation promotes immune evasion by the exclusion of immune cell infiltration into the tumor microenvironment. Using an in vivo model, we observed that the knockdown of NSD1 expression in HNSC tumors established in mice confers a decreased infiltration of CD8^+^ T cells compared to control tumors established in the same animals. The ability of a tumor cell-intrinsic driver to modulate the infiltration of immune cells into the tumor microenvironment has been demonstrated in melanoma, where β-catenin signaling has been shown to result in T cell exclusion, apparently through downregulation of the T cell attractant chemokine CCL4 (34). Moreover, PRC2 mediated epigenetic silencing or chemokines, associated with concordant promoter H3K27me3 and DNA hypermethylation, precludes T cell infiltration in ovarian cancer (63). There was a significant reduction in the expression of several key chemokines associated with knocking down NSD1 in HNSC cell lines, indicating that NSD1 contributes to the regulated expression of these genes in the tumor cells. Efforts are underway to elucidate these mechanisms.

HNSC prognosis has shown little improvement in recent decades (4). Immunotherapies such as monoclonal antibodies to PD-1 or PD-L1, which block the PD-1/PD-L1 checkpoint to restore anti-tumor immune response, are beneficial in a subset of HNSC cases, including metastatic or refractory HNSC cases (64). There is a need to identify biomarkers to predict immunotherapy response, particularly as these treatments can cause autoimmune side effects (65).

As the NSD1 subtype is depleted for both CD8^+^ T cells and PD-1 expressing TAMs, the HNSC NSD1 subtype may be particularly resistant to PD-1/PD-L1 checkpoint blockade immunotherapy, especially as immunotherapy response appears to be dependent on the presence of a preexisting immune cell population (66). The mechanism by which NSD1 inactivation mediates immunosuppression remains to be determined. Most likely, NSD1 inactivation causes epigenetic deregulation of regulators of immune infiltration. Many such genes are epigenetically deregulated in the NSD1 subtype, representing a list of candidate immune regulators that may be investigated in future studies. Such immune regulators may include potential drug targets to restore anti-tumor immunity in NSD1 inactivated HNSCs.

Overall, this study reveals that *NSD1* inactivation confers widespread impairment of epigenetic regulation in both HNSC and LUSC, resulting in loss of epigenetic repression of potential oncogenes. In HNSC, NSD1 inactivation decreases immune cell infiltration, perhaps due to epigenetic deregulation of chemokines. Mechanistic studies into the epigenetic function of NSD1 and classification of the pathways deregulated due to NSD1 inactivation may yield insight that could be exploited to develop novel targeted therapies.

## Conflict of interest

The authors declare that they have no conflicts of interest with the contents of this article

---

**Supplementary figure 1:**
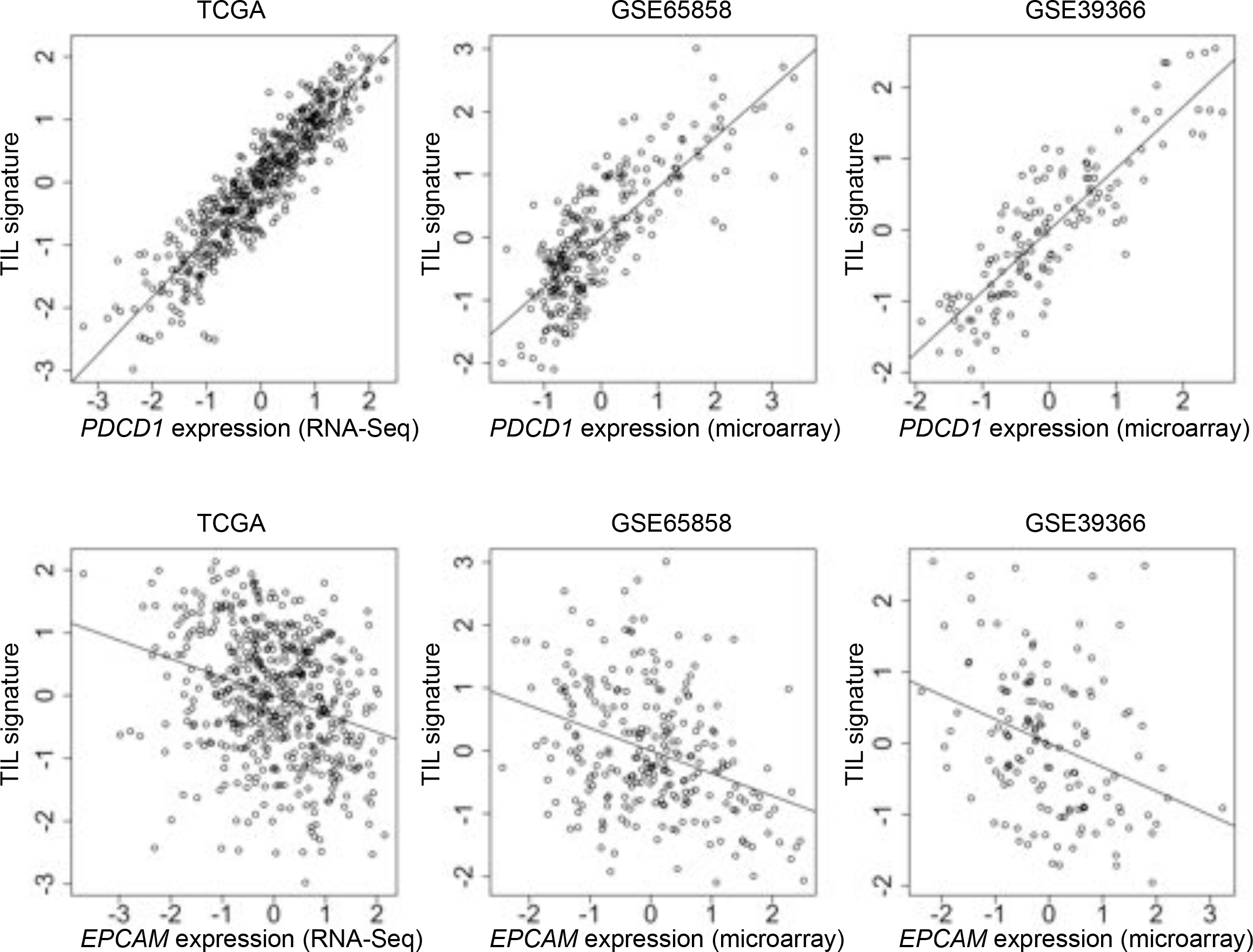
Positive correlations of a T cell signature with expression of PDCD1 expression and inverse correlation with EPCAM (RNA-Seq).

**Supplementary figure 2:**
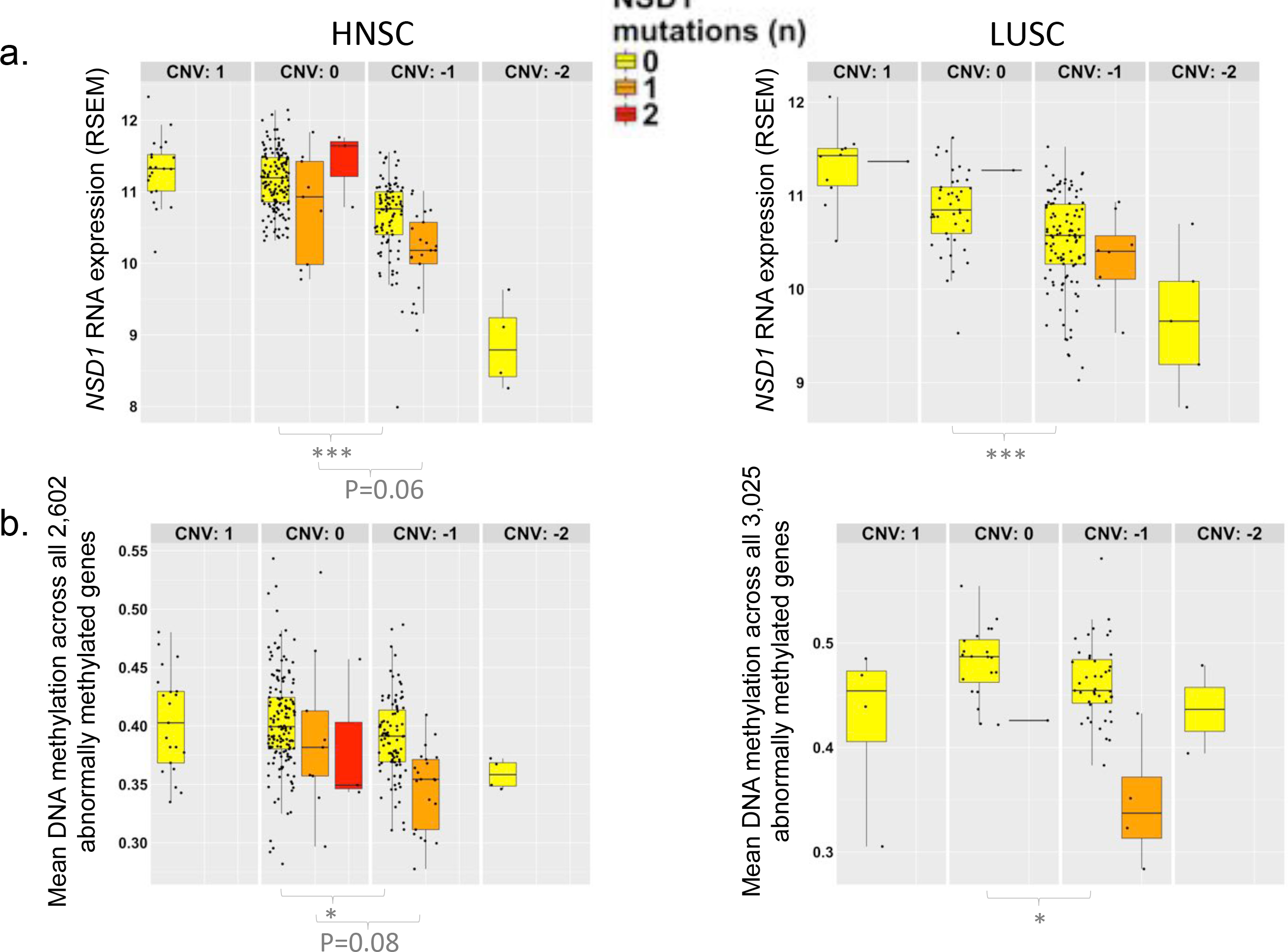
Association of NSD1 inactivating lesions with NSD1 RNA expression and genome-wide abnormal DNA methylation. Levels of a) *NSD1* expression (RNA-Seq) and b) genome-wide DNA methylation, within patient stratified by *NSD1* somatic copy number (Panels) and number of *NSD1* mutations (Colored boxes), indicated for head and neck squamous cell carcinoma (HNSC) and lung squamous cell carcinoma (LUSC) separately. Asterisks indicate statistical significance of difference in *NSD1* expression between indicated groups. ‘Genome-wide abnormal DNA methylation’ represents mean DNA methylation across all abnormally methylated genes, i.e. genes that were either hypermethylated or hypomethylated in tumor relative to normal tissue. *Wilcoxon rank sum test p-value <0.05. ***Wilcoxon rank sum test p-value <0.001.

**Supplementary figure 3:**
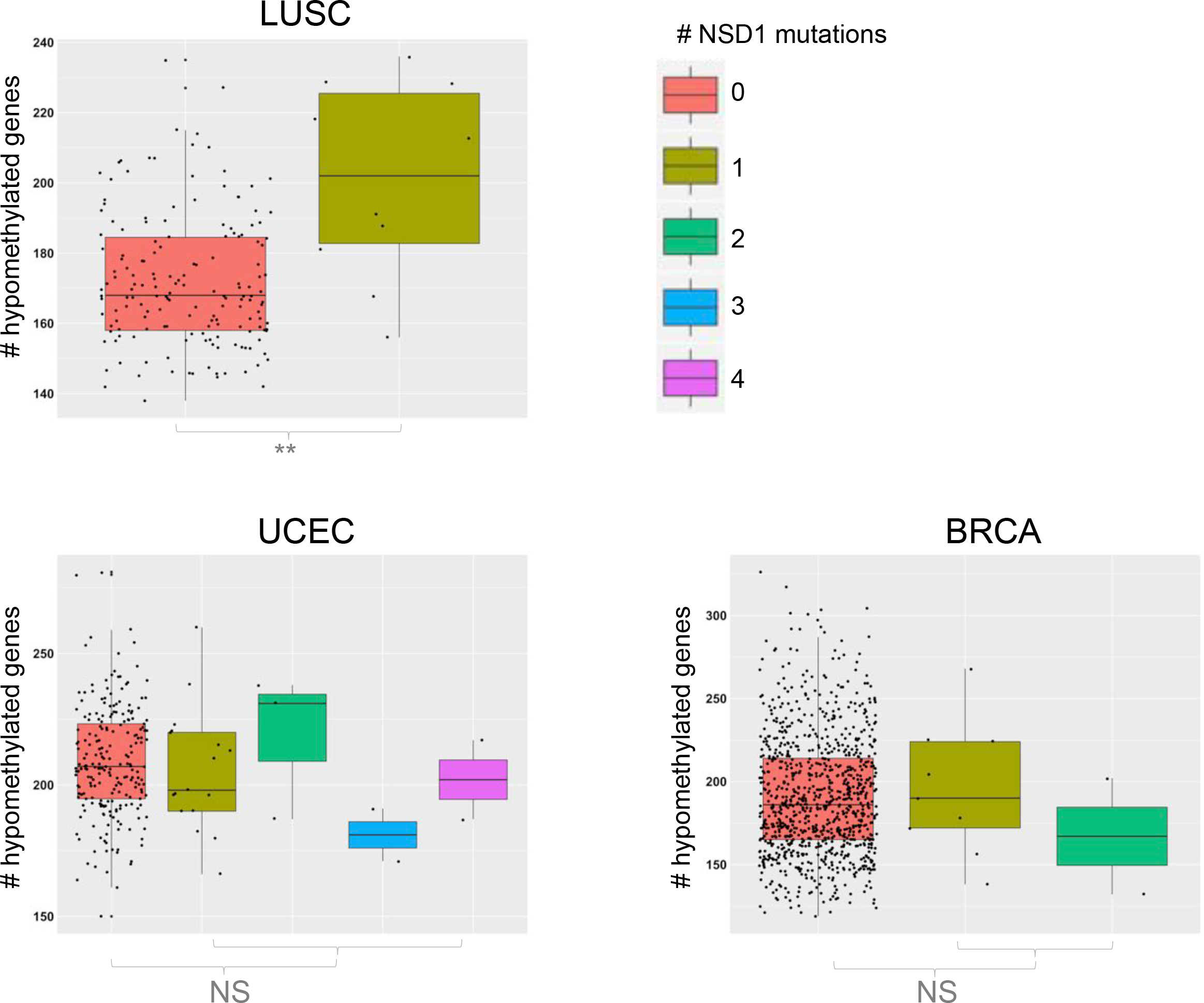
Pan-cancer analysis of DNA hypomethylation associated with NSD1 mutations. Boxplots indicate the number of hypomethylated genes, stratified by *NSD1* mutation status (i.e. number of *NSD1* mutations), for three cancer types (TCGA cancers for which there were at least 10 patients with *NSD1* mutations and DNA methylation data). These include lung squamous cell carcinoma (LUSC, n=10 patients with *NSD1* mutations), uterine corpus endometrial carcinoma (UCEC, n=25 patients with *NSD1* mutations), breast carcinoma (BRCA, n=11 patients with *NSD1* mutations).

**Supplementary figure 4:**
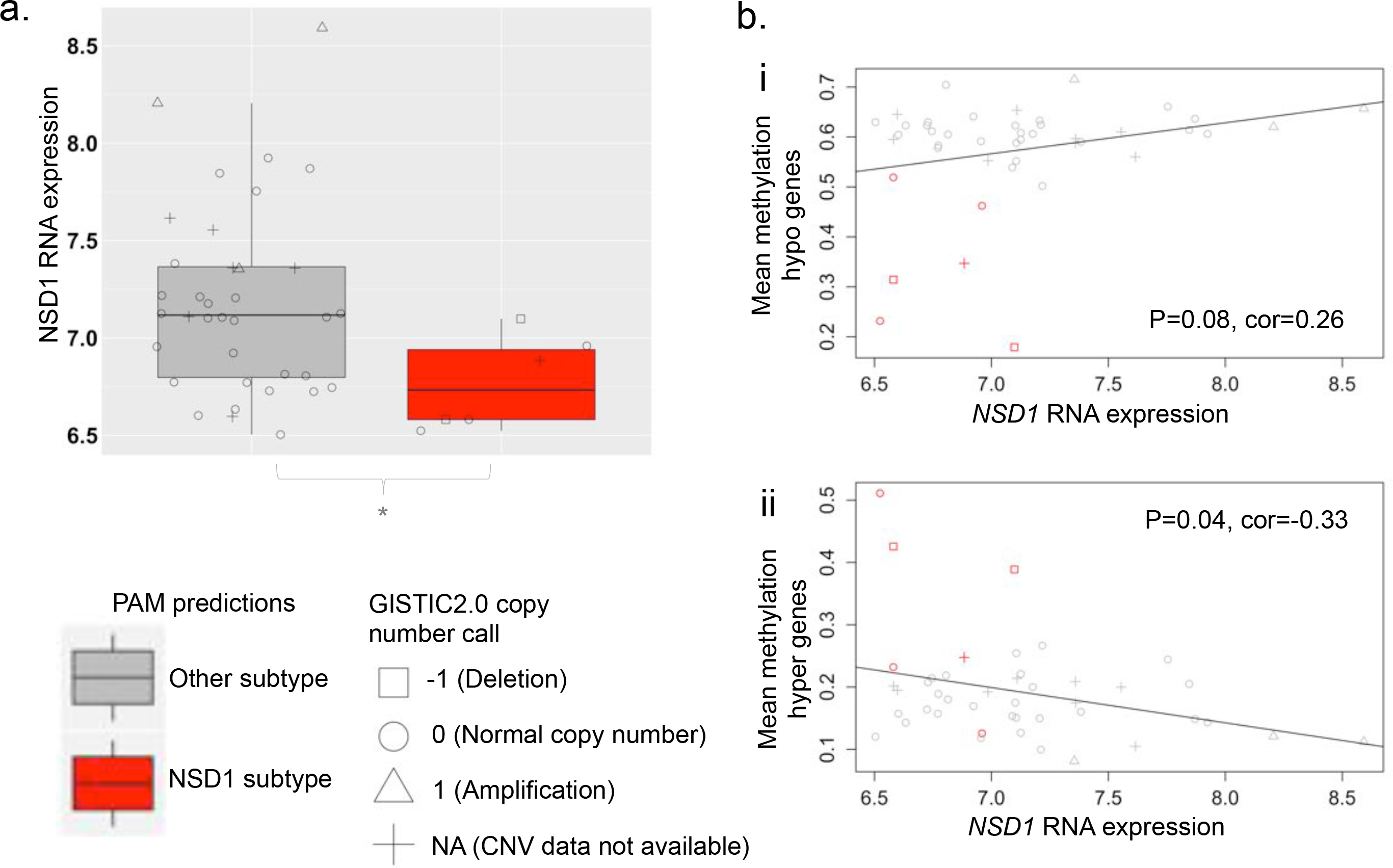
Association of NSD1 RNA expression and copy number with PAM model classes in independent patient cohort. Patients within a validation set of 44 primary HNSCs (GSE33232) were split into two groups based on class prediction for belonging to the NSD1 subtype, using a PAM model that was trained on TCGA DNA methylation data. The asterisk represents the Wilcoxon rank sum test p-value for a difference in mean *NSD1* expression (RNA-Seq) between patients of the predicted NSD1 subtype class (red) and the other class (grey). *P value <0.05 & >0.01. Correlation of *NSD1* expression with DNA methylation of CpG sites that were i) hypomethylated (Supplementary table 1) and ii) hypermethylated (Supplementary table 2) in the TCGA NSD1 subtype, in the GSE33232 cohort. Selected hypomethylated (n=37) and hypermethylated (n=34) CpGs represent those that were within the top 100 genes that were most hypomethylated and hypermethylated genes in the NSD1 subtype relative to other subtypes, in the TCGA cohort. These CpG sites represented those that overlapped between the platforms used to measure DNA methylation in the GSE33232 (Illumina 27k array) and TCGA (Illumina 450k array) cohorts. Patients predicted as belonging to the NSD1 subtype based on the PAM model are highlighted in red. Linear regression P-values and Pearson correlation coefficients (cor) are indicated. *NSD1* copy number calls are, inferred using GISTIC2.0., are indicated by point shapes.

**Supplementary figure 5:**
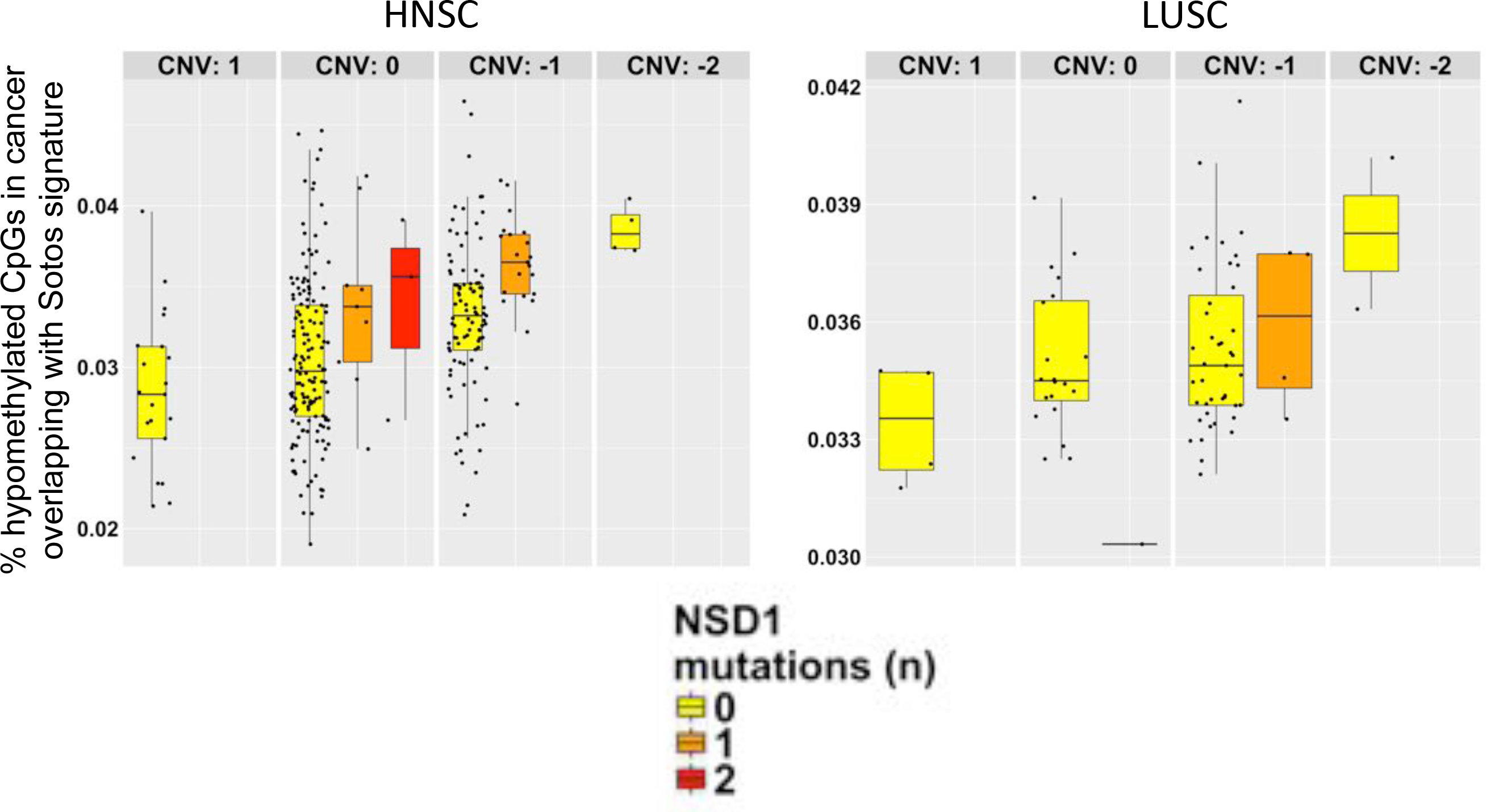
Overlap between aberrant DNA methylation signatures associated with cancer and Sotos syndrome. The percentage of tumor hypomethylated CpG probes (Hypomethylated in tumor relative to normal tissue), that overlapped with 7,038 CpG probes hypomethylated in Sotos syndrome (20) was measured for each cancer patient, representing indices of overall overlap between cancer and Sotos syndrome hypomethylation signatures. *NSD1* mutation count (illustrated by colored boxes) and NSD1 copy number (GISTIC2.0 calls) (Split into panels) are indicated, illustrating the effect of NSD1 inactivation on overlap between cancer and Sotos syndrome signatures.

**Supplementary figure 6:**
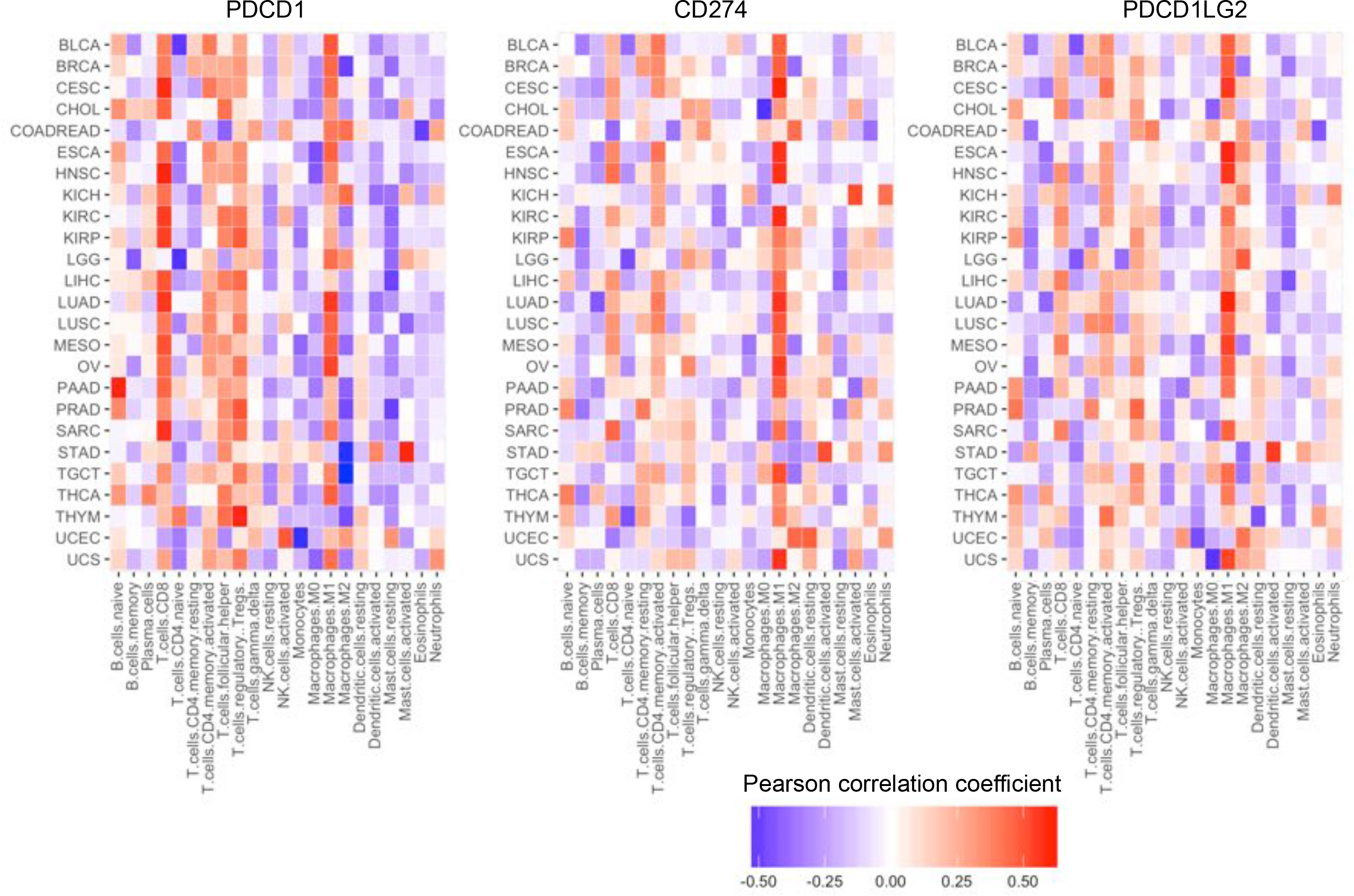
Correlation of PDCD1 (PD-1) expression with inferred levels of tumor associated leukocyte levels in 28 TCGA cancers. Heatmap indicates coefficients for correlation (Pearson) of *PDCD1* RNA expression with levels of CD8+ T cells, M0, M1 and M1 tumor associated macrophages (TAMs), in 28 cancers, using data derived from the TCGA study.

**Supplementary figure 7:**
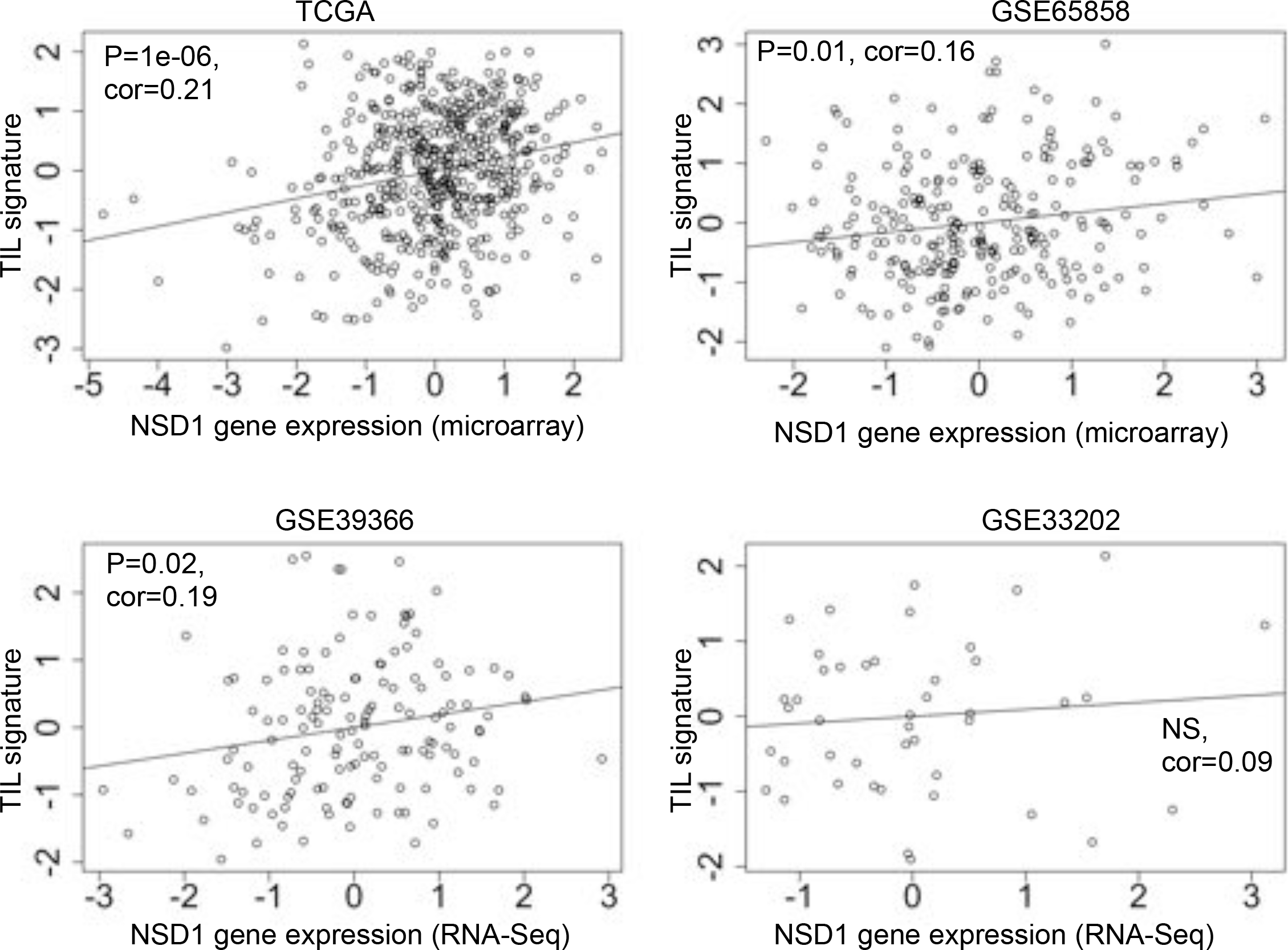
Validation of association of NSD1 RNA expression with a T cell transcript signature in independent patient cohorts. Association of a13 gene T cell transcript signature within the TCGA cohort (Shown for reference), and within gene expression datasets for three independent primary HNSC cohorts, including GSE65858 (n=253) (49), GSE39366 (n=138) and GSE33232 (n=44) (36).

Supplementary table 1: Genes overexpressed/hypomethylated in NSD1 subtypes of HNSC and LUSC

Supplementary table 2: Genes underexpressed/hypermethylated in NSD1 subtypes of HNSC and LUSC

## References

1. Seiwert TY, Zuo Z, Keck MK, Khattri a., Pedamallu CS, Stricker T, et al. Integrative and Comparative Genomic Analysis of HPV-Positive and HPV-Negative Head and Neck Squamous Cell Carcinomas. Clin Cancer Res [Internet]. 2015;21(3):632–41. Available from: http://clincancerres.aacrjournals.org/cgi/doi/10.1158/1078-0432.CCR-13-3310

2. Lawrence MS, Sougnez C, Lichtenstein L, Cibulskis K, Lander E, Gabriel SB, et al. Comprehensive genomic characterization of head and neck squamous cell carcinomas. Nature [Internet]. 2015;517(7536):576–82. Available from: http://www.nature.com/doifinder/10.1038/nature14129

3. Siegel RL, Miller KD, Jemal A. Cancer Statistics, 2015. CA Cancer J Clin. 2015;65(1):5–29.

4. Chuang S-C, Scelo G, Tonita JM, Tamaro S, Jonasson JG, Kliewer E V, et al. Risk of second primary cancer among patients with head and neck cancers: A pooled analysis of 13 cancer registries. Int J Cancer [Internet]. 2008;123(10):2390–6. Available from: http://www.ncbi.nlm.nih.gov/pubmed/18729183

5. Jones S, Stransky N, McCord CL, Cerami E, Lagowski J, Kelly D, et al. Genomic analyses of gynaecologic carcinosarcomas reveal frequent mutations in chromatin remodelling genes. Nat Commun [Internet]. 2014;5:5006. Available from: http://www.ncbi.nlm.nih.gov/pubmed/25233892%5Cnhttp://www.pubmedcentral.nih.gov/articlerender.fcgi?artid=PMC4354107

6. The Cancer Genome Atlas Research Network. Comprehensive genomic characterization of squamous cell lung cancers. Nature [Internet]. 2012;489(7417):519–25. Available from: http://www.pubmedcentral.nih.gov/articlerender.fcgi?artid=3466113&tool=pmcentrez&rendertype=abstract

7. Beltran H, Prandi D, Mosquera JM, Benelli M, Puca L, Cyrta J, et al. Divergent clonal evolution of castration-resistant neuroendocrine prostate cancer. Nat Med [Internet]. 2016;22(3):298–305. Available from: http://www.ncbi.nlm.nih.gov/pubmed/26855148

8. Lucio-Eterovic AK, Carpenter PB. An open and shut case for the role of NSD proteins as oncogenes. Transcription [Internet]. 2011;2(4): 158–61. Available from: http://www.pubmedcentral.nih.gov/articlerender.fcgi?artid=3173681&tool=pmcentrez&rendertype=abstract

9. Cancer Genome Atlas Research Network. Comprehensive molecular characterization of clear cell renal cell carcinoma. Nature [Internet]. 2013;499(7456):43–9. Available from: http://www.ncbi.nlm.nih.gov/pubmed/23792563

10. Berdasco M, Ropero S, Setien F, Fraga MF, Lapunzina P, Losson R, et al. Epigenetic inactivation of the Sotos overgrowth syndrome gene histone methyltransferase NSD1 in human neuroblastoma and glioma. Proc Natl Acad Sci U S A [Internet]. 2009;106(51):21830–5. Available from: http://www.pnas.org.ezp-prod1.hul.harvard.edu/content/106/51/21830.long

11. Shiba N, Ichikawa H, Taki T, Park M, Jo A, Mitani S, et al. NUP98-NSD1 gene fusion and its related gene expression signature are strongly associated with a poor prognosis in pediatric acute myeloid leukemia. Genes Chromosomes Cancer [Internet]. 2013;52(7):683–93. Available from: http://www.ncbi.nlm.nih.gov/pubmed/23630019

12. Quintana RM, Dupuy AJ, Bravo A, Casanova ML, Alameda JP, Page A, et al. A transposon-based analysis of gene mutations related to skin cancer development. J Invest Dermatol [Internet]. 2013;133(1):239–48. Available from: http://www.ncbi.nlm.nih.gov/pubmed/22832494

13. Douglas J, Hanks S, Temple IK, Davies S, Murray A, Upadhyaya M, et al. NSD1 mutations are the major cause of Sotos syndrome and occur in some cases of Weaver syndrome but are rare in other overgrowth phenotypes. Am J Hum Genet. 2003;72(1):132–43.

14. Kurotaki N, Imaizumi K, Harada N, Masuno M, Kondoh T, Nagai T, et al. Haploinsufficiency of NSD1 causes Sotos syndrome. Nat Genet. 2002;30(4):365–6.

15. Baujat G, Rio M, Rossignol S, Sanlaville D, Lyonnet S, Le Merrer M, et al. Paradoxical NSD1 mutations in Beckwith-Wiedemann syndrome and 11p15 anomalies in Sotos syndrome. Am J Hum Genet. 2004;74(4):715–20.

16. Feinberg AP, Koldobskiy MA, Gondor A. Epigenetic modulators, modifiers and mediators in cancer aetiology and progression. Nat Rev Genet [Internet]. 2016;17(5):284–99. Available from: http://www.nature.com/doifinder/10.1038/nrg.2016.13%5Cnhttp://www.ncbi.nlm.nih.gov/pubmed/26972587

17. Qiao Q, Li Y, Chen Z, Wang M, Reinberg D, Xu RM. The structure of NSD1 reveals an autoregulatory mechanism underlying histone H3K36 methylation. J Biol Chem. 2011;286(10):8361–8.

18. Tatton-Brown K, Rahman N. The NSD1 and EZH2 overgrowth genes, similarities and differences. Am J Med Genet Part C Semin Med Genet. 2013;163(2):86–91.

19. Papillon-Cavanagh S, Lu C, Gayden T, Mikael LG, Bechet D, Karamboulas C, et al. Impaired H3K36 methylation defines a subset of head and neck squamous cell carcinomas. Nat Genet [Internet]. Nature Publishing Group, a division of Macmillan Publishers Limited. All Rights Reserved.; 2017 Feb;49(2):180–5. Available from: http://dx.doi.org/10.1038/ng.3757

20. Choufani S, Cytrynbaum C, Chung BHY, Turinsky AL, Grafodatskaya D, Chen YA, et al. NSD1 mutations generate a genome-wide DNA methylation signature. Nat Commun [Internet]. 2015;6:10207. Available from: http://www.nature.com/doifinder/10.1038/ncomms10207

21. Gevaert O, Tibshirani R, Plevritis SK. Pancancer analysis of DNA methylation-driven genes using MethylMix. Genome Biol. 2015;1–13.

22. Brennan K, Koenig JL, Gentles AJ, Sunwoo JB, Gevaert O. Identification of an atypical etiological head and neck squamous carcinoma subtype featuring the CpG island methylator phenotype. EBioMedicine [Internet]. Elsevier B.V; 2017; Available from: http://linkinghub.elsevier.com/retrieve/pii/S2352396417300841

23. Daud AI, Loo K, Pauli ML, Sanchez-Rodriguez R, Sandoval PM, Taravati K, et al. Tumor immune profiling predicts response to anti-PD-1 therapy in human melanoma. J Clin Invest. 2016;126(9):3447–52.

24. Samur MK. RTCGAToolbox: A New Tool for Exporting TCGA firehose data. PLoS One. 2014;9(9).

25. Lawrence MS, Stojanov P, Polak P, Kryukov G V, Cibulskis K, Sivachenko A, et al. Mutational heterogeneity in cancer and the search for new cancer-associated genes. Nature [Internet]. 2013;499(7457):214–8. Available from: http://dx.doi.org/10.1038/nature12213

26. Troyanskaya O, Cantor M, Sherlock G, Brown P, Hastie T, Tibshirani R, et al. Missing value estimation methods for DNA microarrays. Bioinformatics. 2001;17(6):520–5.

27. Johnson WE, Li C, Rabinovic A. Adjusting batch effects in microarray expression data using empirical Bayes methods. Biostatistics. 2007;8(1):118–27.

28. Gevaert O. MethylMix: an R package for identifying DNA methylation driven genes. Bioinformatics. 2015 Jan;btv020.

29. Wilkerson MD, Hayes DN. ConsensusClusterPlus: A class discovery tool with confidence assessments and item tracking. Bioinformatics. 2010;26(12):1572–3.

30. Tusher VG, Tibshirani R, Chu G. Significance analysis of microarrays applied to the ionizing radiation response. Proc Natl Acad Sci U S A. 2001;98(9):5116–21.

31. Tibshirani R, Hastie T, Narasimhan B, Chu G. Diagnosis of multiple cancer types by shrunken centroids of gene expression. Proc Natl Acad Sci U S A. 2002;99(10):6567–72.

32. Newman AM, Liu CL, Green MR, Gentles AJ, Feng W, Xu Y, et al. Robust enumeration of cell subsets from tissue expression profiles. Nat Methods [Internet]. 2015;(MAY 2014): 1–10. Available from: http://www.nature.com/doifinder/10.1038/nmeth.3337%5Cnhttp://www.ncbi.nlm.nih.gov/pubmed/25822800

33. Gentles AJ, Newman AM, Liu CL, Bratman S V, Feng W, Kim D, et al. The prognostic landscape of genes and infiltrating immune cells across human cancers. Nat Med [Internet]. 2015;21(8):938–45. Available from: http://www.ncbi.nlm.nih.gov/pubmed/26193342

34. Spranger S, Bao R, Gajewski TF. Melanoma-intrinsic β-catenin signalling prevents antitumour immunity. Nature [Internet]. 2015; Available from: http://www.nature.com/doifinder/10.1038/nature14404

35. Yoshihara K, Shahmoradgoli M, Martínez E, Vegesna R, Kim H, Torres-Garcia W, et al. Inferring tumour purity and stromal and immune cell admixture from expression data. Nat Commun [Internet]. 2013;4:2612. Available from: http://www.pubmedcentral.nih.gov/articlerender.fcgi?artid=3826632&tool=pmcentrez&rendertype=abstract

36. Fertig EJ, Markovic A, Danilova L V., Gaykalova DA, Cope L, Chung CH, et al. Preferential activation of the hedgehog pathway by epigenetic modulations in HPV negative HNSCC identified with meta-pathway analysis. PLoS One. 2013;8(11).

37. Korn JM, Kuruvilla FG, McCarroll S a, Wysoker A, Nemesh J, Cawley S, et al. Integrated genotype calling and association analysis of SNPs, common copy number polymorphisms and rare CNVs. Nat Genet [Internet]. 2008;40(10):1253–60. Available from: http://www.ncbi.nlm.nih.gov/pubmed/18776909

38. Mermel CH, Schumacher SE, Hill B, Meyerson ML, Beroukhim R, Getz G. GISTIC2.0 facilitates sensitive and confident localization of the targets of focal somatic copy-number alteration in human cancers. Genome Biol [Internet]. 2011;12(4):R41. Available from: http://www.ncbi.nlm.nih.gov/pubmed/21527027

39. Walter V, Yin X, Wilkerson MD, Cabanski CR, Zhao N, Du Y, et al. Molecular subtypes in head and neck cancer exhibit distinct patterns of chromosomal gain and loss of canonical cancer genes. PLoS One [Internet]. 2013;8(2):e56823. Available from: http://www.pubmedcentral.nih.gov/articlerender.fcgi?artid=3579892&tool=pmcentrez&rendertype=abstract

40. Qu X, Liu J, Zhong X, Li X, Zhang Q. PIWIL2 promotes progression of non-small cell lung cancer by inducing CDK2 and Cyclin A expression. J Transl Med [Internet].2015; 13(1):301. Available from: http://www.translational-medicine.com/content/13/1/301%5Cnhttp://translational-medicine.biomedcentral.com/articles/10.1186/s12967-015-0666-y

41. Lee JH, Schütte D, Wulf G, Füzesi L, Radzun HJ, Schweyer S, et al. Stem-cell protein Piwil2 is widely expressed in tumors and inhibits apoptosis through activation of Stat3/Bcl-XL pathway. Hum Mol Genet. 2006;15(2):201–11.

42. Ng RK, Dean W, Dawson C, Lucifero D, Madeja Z, Reik W, et al. Epigenetic restriction of embryonic cell lineage fate by methylation of Elf5. Nat Cell Biol [Internet]. 2008;10(11): 1280–1290. Available from: http://www.nature.com/ncb/journal/vaop/ncurrent/full/ncb1786.html

43. Takemoto T, Uchikawa M, Yoshida M, Bell DM, Lovell-Badge R, Papaioannou VE, et al. Tbx6-dependent Sox2 regulation determines neural or mesodermal fate in axial stem cells. Nature [Internet]. 2011;470(7334):394–8. Available from: http://www.pubmedcentral.nih.gov/articlerender.fcgi?artid=3042233&tool=pmcentrez&rendertype=abstract

44. Chiu WT, Charney Le R, Blitz IL, Fish MB, Li Y, Biesinger J, et al. Genome-wide view of TGFβ/Foxh1 regulation of the early mesendoderm program. Development [Internet]. 2014;141(23):4537–47. Available from: http://www.ncbi.nlm.nih.gov/pubmed/25359723

45. Gordon SR, Maute RL, Dulken BW, Hutter G, George BM, McCracken MN, et al. PD-1 expression by tumour-associated macrophages inhibits phagocytosis and tumour immunity. Nature [Internet]. Nature Publishing Group; 2017;545(7655):495–9. Available from: http://www.nature.com/doifinder/10.1038/nature22396

46. Schalper K, Carvajal-Hausdorf D, McLaughlin J, Velcheti V, Chen L, Sanmamed M, et al. Clinical significance of PD-L1 protein expression on tumor-associated macrophages in lung cancer. J Immunother Cancer [Internet]. BioMed Central Ltd; 2015;3(Suppl 2):P415. Available from: http://jitc.biomedcentral.com/articles/10.1186/2051-1426-3-S2-P415

47. Haderk F, Schulz R, Iskar M, Cid LL, Worst T, Willmund K V, et al. Tumor-derived exosomes modulate PD-L1 expression in monocytes. Sci Immunol [Internet]. 2017;2(July):1–12. Available from: http://immunology.sciencemag.org/content/immunology/2/13/eaah5509.full.pdf

48. Hartley G, Regan D, Guth A, Dow S. Regulation of PD-L1 expression on murine tumor-associated monocytes and macrophages by locally produced TNF-α. Cancer Immunol Immunother [Internet]. 2017 Apr;66(4):523–35. Available from: https://doi.org/10.1007/s00262-017-1955-5

49. Gunnar Wichmann1, 2*, Maciej Rosolowski, 2, 3* KK, Kreuz3 M, Boehm1 A, Reiche1 A, Ulrike Scharrer1, 2 DH, Bertolini6 J, et al. The role of HPV RNA transcription, immune response-related gene expression and disruptive TP53 mutations in diagnostic and prognostic profiling of head and neck cancer. 2015;0.

50. You JS, Jones PA. Cancer Genetics and Epigenetics: Two Sides of the Same Coin? Vol.22, Cancer Cell. 2012. p. 9–20.

51. Weisenberger DJ. Characterizing DNA methylation alterations from the cancer genome atlas. Vol. 124, Journal of Clinical Investigation. 2014. p. 17–23.

52. Tiedemann RL, Hlady RA, Hanavan PD, Lake DF, Tibes R, Lee J, et al. Dynamic reprogramming of DNA methylation in SETD2-deregulated renal cell carcinoma. Oncotarget [Internet]. 2016;7(2):1927–46. Available from: http://www.ncbi.nlm.nih.gov/pubmed/26646321

53. Baubec T, Colombo DF, Wirbelauer C, Schmidt J, Burger L, Krebs AR, et al. Genomic profiling of DNA methyltransferases reveals a role for DNMT3B in genic methylation. Nature [Internet]. 2015;520(7546):243–7. Available from: http://www.ncbi.nlm.nih.gov/pubmed/25607372%5Cnhttp://dx.doi.org/10.1038/nature14176%5Cnhttp://www.nature.com/doifinder/10.1038/nature14176

54. Dhayalan A, Rajavelu A, Rathert P, Tamas R, Jurkowska RZ, Ragozin S, et al. The Dnmt3a PWWP domain reads histone 3 lysine 36 trimethylation and guides DNA methylation. J Biol Chem. 2010;285(34):26114–20.

55. Lucio-Eterovic AK, Singh MM, Gardner JE, Veerappan CS, Rice JC, Carpenter PB. Role for the nuclear receptor-binding SET domain protein 1 (NSD1) methyltransferase in coordinating lysine 36 methylation at histone 3 with RNA polymerase II function. Proc Natl Acad Sci U S A. 2010;107(39):16952–7.

56. Wagner EJ, Carpenter PB. Understanding the language of Lys36 methylation at histone H3. Nat Rev Mol Cell Biol [Internet]. 2012; 13(2): 115–26. Available from: http://dx.doi.org/10.1038/nrm3274%5Cnhttp://www.pubmedcentral.nih.gov/articlerender.fcgi?artid=3969746&tool=pmcentrez&rendertype=abstract

57. Sherwood AM, Emerson RO, Scherer D, Habermann N, Buck K, Staffa J, et al. Tumor-infiltrating lymphocytes in colorectal tumors display a diversity of T cell receptor sequences that differ from the T cells in adjacent mucosal tissue. Cancer Immunol Immunother. 2013;62(9):1453–61.

58. Loi S, Michiels S, Salgado R, Sirtaine N, Jose V, Fumagalli D, et al. Tumor infiltrating lymphocytes are prognostic in triple negative breast cancer and predictive for trastuzumab benefit in early breast cancer: results from the FinHER trial. Ann Oncol [Internet]. 2014;25(8):1544–50. Available from: http://www.ncbi.nlm.nih.gov/pubmed/24608200

59. Nielsen JS, Sahota R a., Milne K, Kost SE, Nesslinger NJ, Watson PH, et al. CD20^+^ tumor-infiltrating lymphocytes have an atypical CD27 - memory phenotype and together with CD8^+^ T cells promote favorable prognosis in ovarian cancer. Clin Cancer Res. 2012;18(12):3281–92.

60. Yaguchi T, Goto Y, Kido K, Mochimaru H, Sakurai T, Tsukamoto N, et al. Immune suppression and resistance mediated by constitutive activation of Wnt/β-catenin signaling in human melanoma cells. J Immunol [Internet]. 2012;189(5):2110–7. Available from: http://www.ncbi.nlm.nih.gov/pubmed/22815287

61. Gentles AJ, Newman AM, Liu CL, Bratman S V, Feng W, Kim D, et al. The prognostic landscape of genes and infiltrating immune cells across human cancers. Nat Med [Internet]. Nature Publishing Group; 2015;21(8): 1–12. Available from: http://www.nature.com/doifinder/10.1038/nm.3909

62. Badoual C, Hans S, Rodriguez J, Peyrard S, Klein C, Agueznay NEH, et al. Prognostic value of tumor-infiltrating CD4^+^ T-cell subpopulations in head and neck cancers. Clin Cancer Res. 2006;12(2):465–72.

63. Peng D, Kryczek I, Nagarsheth N, Zhao L, Wei S, Wang W, et al. Epigenetic silencing of TH1-type chemokines shapes tumour immunity and immunotherapy. Nature [Internet]. 2015;527(7577):1–16. Available from: http://www.nature.com/doifinder/10.1038/nature15520%5Cn http://www.ncbi.nlm.nih.gov/pubmed/26503055

64. Seiwert TY, Burtness B, Mehra R, Weiss J, Berger R, Eder JP, et al. Safety and clinical activity of pembrolizumab for treatment of recurrent or metastatic squamous cell carcinoma of the head and neck (KEYNOTE-012): an open-label, multicentre, phase 1b trial. Lancet Oncol [Internet]. Elsevier; 2017 Jul 3;17(7):956–65. Available from: http://dx.doi.org/10.1016/S1470-2045(16)30066-3

65. Naidoo J, Page DB, Li BT, Connell LC, Schindler K, Lacouture ME, et al. Toxicities of the anti-PD-1 and anti-PD-L1 immune checkpoint antibodies. Ann Oncol. 2015;26(12):2375–91.

66. Badoual C, Hans S, Merillon N, Van Ryswick C, Ravel P, Benhamouda N, et al. PD-1-expressing tumor-infiltrating T cells are a favorable prognostic biomarker in HPV associated head and neck cancer. Cancer Res [Internet]. 2012;128–38. Available from: http://www.ncbi.nlm.nih.gov/pubmed/23135914

67. Hoadley KA, Yau C, Wolf DM, Cherniack AD, Tamborero D, Ng S, et al. Multiplatform analysis of 12 cancer types reveals molecular classification within and across tissues of origin. Cell. 2014;158(4):929–44.

68. Liu Z, Zhang S. Tumor characterization and stratification by integrated molecular profiles reveals essential pan-cancer features. BMC Genomics [Internet]. 2015;16(1):503. Available from: http://www.biomedcentral.com/1471-2164/16/503

69. Feinberg AP, Vogelstein B. Hypomethylation distinguishes genes of some human cancers from their normal counterparts. Nature. 1983;301(5895):89–92.

70. Ehrlich M. DNA hypomethylation in cancer cells. Epigenomics [Internet]. 2009;1(2):239–59. Available from: http://www.futuremedicine.com/doi/abs/10.2217/epi.09.33

71. Van Tongelen A, Loriot A, De Smet C. Oncogenic roles of DNA hypomethylation through the activation of cancer-germline genes. Cancer Lett [Internet]. 2017;(March):1–8. Available from: http://www.sciencedirect.com/science/article/pii/S0304383517302033

72. Cannuyer J, Van Tongelen A, Loriot A, De Smet C. A gene expression signature identifying transient DNMT1 depletion as a causal factor of cancer-germline gene activation in melanoma. Clin Epigenetics [Internet]. 2015;7:114. Available from: http://www.pubmedcentral.nih.gov/articlerender.fcgi?artid=4620642&tool=pmcentrez&rendertype=abstract

73. Lee JH, Jung C, Javadian-Elyaderani P, Schweyer S, Schutte D, Shoukier M, et al. Pathways of proliferation and antiapoptosis driven in breast cancer stem cells by stem cell protein piwil2. Cancer Res [Internet]. 2010;70(11):4569–79. Available from: http://www.ncbi.nlm.nih.gov/pubmed/20460541

74. Gallego-Ortega D, Ledger A, Roden DL, Law AMK, Magenau A, Kikhtyak Z, et al. ELF5 Drives Lung Metastasis in Luminal Breast Cancer through Recruitment of Gr1^+^ CD11b^+^ Myeloid-Derived Suppressor Cells. PLoS Biol. 2015; 13(12).

75. Kalyuga M, Gallego-Ortega D, Lee HJ, Roden DL, Cowley MJ, Caldon CE, et al. ELF5 Suppresses Estrogen Sensitivity and Underpins the Acquisition of Antiestrogen Resistance in Luminal Breast Cancer. PLoS Biol. 2012;10(12).

76. Gibson WT, Hood RL, Zhan SH, Bulman DE, Fejes AP, Moore R, et al. Mutations in EZH2 cause weaver syndrome. Am J Hum Genet. 2012;90(1):110–8.

77. Luscan A, Laurendeau I, Malan V, Francannet C, Odent S, Giuliano F, et al. Mutations in SETD2 cause a novel overgrowth condition. J Med Genet [Internet]. 2014;51(8):512–7. Available from: http://www.ncbi.nlm.nih.gov/pubmed/24852293

78. Nimura K, Ura K, Shiratori H, Ikawa M, Okabe M, Schwartz RJ, et al. A histone H3 lysine 36 trimethyltransferase links Nkx2-5 to Wolf-Hirschhorn syndrome. Nature [Internet]. 2009;460(7252):287–91. Available from: http://www.nature.com/doifinder/10.1038/nature08086

79. Hock H. A complex Polycomb issue: The two faces of EZH2 in cancer. Genes Dev. 2012;26(8):751–5.

80. Gartia-Carpizo V, Sarmentero J, Han B, Graña O, Ruiz-Llorente S, Pisano DG, et al. NSD2 contributes to oncogenic RAS-driven transcription in lung cancer cells through long-range epigenetic activation. Sci Rep [Internet]. 2016;6(August):32952. Available from: http://www.nature.com/articles/srep32952

